# *In vivo* human embryonic spinal cord atlas validates stem cell–derived human dorsal interneurons and reveals ASD spinal signatures

**DOI:** 10.64898/2025.12.22.696129

**Authors:** Sandeep Gupta, Eric Heinrichs, Cristian Rodriguez, Emily Friedman, Salena Gallardo, Talin Dermirjie, Teny Panosian, Keith Phan, Anooshik Tahmasian, Yahir Verdin, Samantha J. Butler

**Affiliations:** Department of Neurobiology, David Geffen School of Medicine, University of California, Los Angeles, Los Angeles, CA 90095; Department of Cell Biology, University of Alberta, Edmonton, Canada; Molecular Biology Interdepartmental Graduate Program, University of California, Los Angeles, Los Angeles, CA 90095; CIRM Bridges to Research Program, California State University, Northridge, CA 91330; Eli and Edythe Broad Center of Regenerative Medicine and Stem Cell Research, University of California, Los Angeles, Los Angeles, CA 90095; Intellectual & Developmental Disabilities Research Center, University of California, Los Angeles, Los Angeles, CA 90095

**Keywords:** human, stem cells, spinal cord, sensory, dorsal interneurons, reference atlas, directed differentiation protocol, neuromesodermal progenitors, BMP4, GDF11, HOX genes, posteriorization

## Abstract

Restoring somatosensory function after spinal cord injury (SCI) faces fundamental challenges: neuronal subtypes must match both axial position and circuit identity, yet the developmental patterning of human dorsal spinal interneurons (dIs) remains incompletely defined. Here, we integrate six single-cell transcriptomics datasets derived from human embryonic spinal cord tissue spanning gestational weeks 4-25 to generate a reference atlas of early human somatosensory circuit development. The atlas reveals molecular signatures underlying expansion and specialization of dI4 and dI5 interneuron populations associated with mechanosensory and nociceptive processing. Guided by this resource, we established a neuromesodermal progenitor–based differentiation approach that generates dorsal interneurons spanning anterior–posterior identities. Comparison of *in vivo* and *in vitro* dI4/dI5 subclasses identified conserved gene networks associated with sensory modalities and revealed enrichment of autism spectrum disorder–associated genes within mechanosensory interneuron populations. Together, these findings clarify how human dorsal spinal interneuron diversity is established.

## Introduction

Spinal cord injury (SCI) is a debilitating condition lacking curative therapies^1^. Although loss of motor function is the most widely recognized consequence of SCI, disruption of somatosensory perception - including pain, touch, and proprioception - is equally detrimental for patients’ quality of life^2, 3^. Compared to the recovery of motor function, recovery of sensory function is highly variable, and many individuals with severe injuries experience persistent deficits in sensory perception^4^. Thus, the restoration of somatosensory circuitry represents a critical but understudied challenge in regenerative neuroscience.

Cell replacement strategies offer a promising approach to repair damaged spinal circuits^5, 6^. Advances in stem cell biology now permit the generation of neuronal subtypes capable of integrating into host tissue^7–9^. However, successful circuit reconstruction in the spinal cord is likely to require precise matching of neuronal identity to anatomical position along the anterior–posterior axis. As the spinal cord is organized into segmental domains that connect distinct peripheral organs and tissues^10^, restoring function requires generating neuronal subtypes with matching regional identities. For example, restoration of bladder and bowel function depends on reconstruction of lumbosacral circuits that regulate autonomic control^11^. Achieving such specificity requires differentiation strategies that reproduce developmental programs governing spinal cord regionalization.

The signals that determine regional identity have been well characterized in the developing spinal cord^12, 13^. Anterior/posterior identity is established by the expression of *HOX* transcription factors^13–15^. *HOX* genes are organized into four chromosomal clusters (*HOXA*, *HOXB*, *HOXC,* and *HOXD)* and activated in a temporally progressive manner^16–18^: anterior identities are specified by 3’ HOX paralogs while more posterior identities emerge through later expression of the 5’ HOX genes^18^. In parallel, signaling centers within the neural tube establish dorsal-ventral patterning by secreting growth factors^19^. In the dorsal spinal cord, six dorsal progenitor (dP) populations generate six classes of dorsal interneurons (dI1-dI6) that process somatosensory inputs including touch, pain, temperature, and proprioception^20^. Ventral progenitor domains produce motor neurons (MNs) and ventral interneurons that coordinate locomotor output^21, 22^. Together, these patterning mechanisms generate the cellular diversity required for sensorimotor circuit assembly.

Directed differentiation protocols have successfully generated spinal MNs from pluripotent stem cells that achieved functional integration in experimental SCI models^23–25^. However, the restoration of somatosensory circuitry will also require the robust generation of dIs spanning multiple axial levels. Early differentiation protocols recapitulated aspects of spinal patterning by inducing anterior neuroepithelium followed by posteriorization toward spinal identities^26–29^. While these approaches generate MNs and a subset of dI populations with brachial-thoracic identities, they do not efficiently produce neurons corresponding to the most posterior spinal regions^26, 28^. *In vivo*, much of the posterior spinal cord arises from bipotential neuromesodermal progenitors (NMPs)^30^, which give rise to the lumbar-sacral circuits essential for bladder and bowel control^30, 31^. Importantly, NMP-based differentiation strategies show improved outcomes; a recent mouse embryonic stem cell (ESC) protocol that derived dIs through an NMP intermediate produced the full complement of dIs that transcriptionally and functionally resemble their endogenous counterparts^32, 33^. However, comparable approaches for generating human dIs across the full anterior-posterior axis have yet to be developed.

A major barrier to evaluating stem cell-derived spinal neurons is the lack of a comprehensive human developmental reference atlas. Directed differentiation protocols typically produce heterogenous neuronal populations, and the validation of cellular identity requires comparison to endogenous developmental trajectories^28^. Advances in single-cell transcriptomics and sophisticated computational methods have enabled the integration of large-scale datasets to reconstruct developmental lineages across tissues^34, 35^. While such approaches have been applied broadly in other organ systems, the application to human embryonic spinal cord development has been more limited. Thus, the existing datasets for the human embryonic spinal cord provide an incomplete view of neuronal diversity and lineage relationships, limiting the ability to benchmark *in vitro*-derived cell types or identify previously unrecognized neuronal subclasses. A comprehensive reference atlas of the developing human spinal cord would therefore provide an important resource for defining neuronal identity, mapping developmental trajectories, and guiding rational design of differentiation strategies for regenerative applications.

Here, we integrated six publicly available single-cell datasets^18, 36–40^ to generate a transcriptomic reference atlas of the developing human spinal cord spanning gestational weeks 4-25. The atlas reveals extensive diversification of dI populations, including a marked expansion in the dI4 and dI5 subclasses associated with somatosensory processing of pain, touch, and itch^10^. Using this resource as a benchmark, we developed a human NMP-based differentiation protocol designed to generate dIs across the anterior-posterior axis. By modulating NMP culture duration and exposure to the posteriorizing factor GDF11^41–43^, we produced interneuron populations that align with endogenous developmental trajectories and exhibit molecular signatures consistent with somatosensory circuit specialization. Comparative analysis of *in vivo* and *in vitro* dI4/dI5 subclasses reveal conserved sensory-associated gene networks, including enrichment of genes associated with autism spectrum disorder (ASD), suggesting that molecular programs linked to sensory phenotypes emerge during early spinal cord development. Taken together, these findings provide an experimental paradigm for investigating the organization of human somatosensory circuitry and establish principles for generating regionally specified spinal interneurons for disease modeling and regenerative strategies.

## Results

### Creation of a human fetal single-cell atlas resolves the developmental trajectories of spinal cord cell types

To establish a reference dataset for human spinal cord development, we integrated transcriptomic datasets spanning fetal development to generate a harmonized single-cell atlas of the human spinal cord (Fig. 1b). We compiled 192 publicly available single cell (sc) -and single-nucleus (sn) RNA-Seq samples from six studies^18, 36–40^, yielding an integrated dataset of approximately 1.7 million cells spanning gestational weeks 4-25 (Extended Data Fig. 1a-c). Datasets were preprocessed using Seurat^44^ and integrated using the deep generative modeling program SCVI^45^ to enable robust cross-study alignment while preserving biological structure (Fig. 1a) (see Materials and Methods for a complete description of the computational pipeline). The atlas can be viewed online here: https://cells.ucsc.edu/?ds=spinal-cord-asd. (Extended Data Fig. 1b).

**Figure 1:**
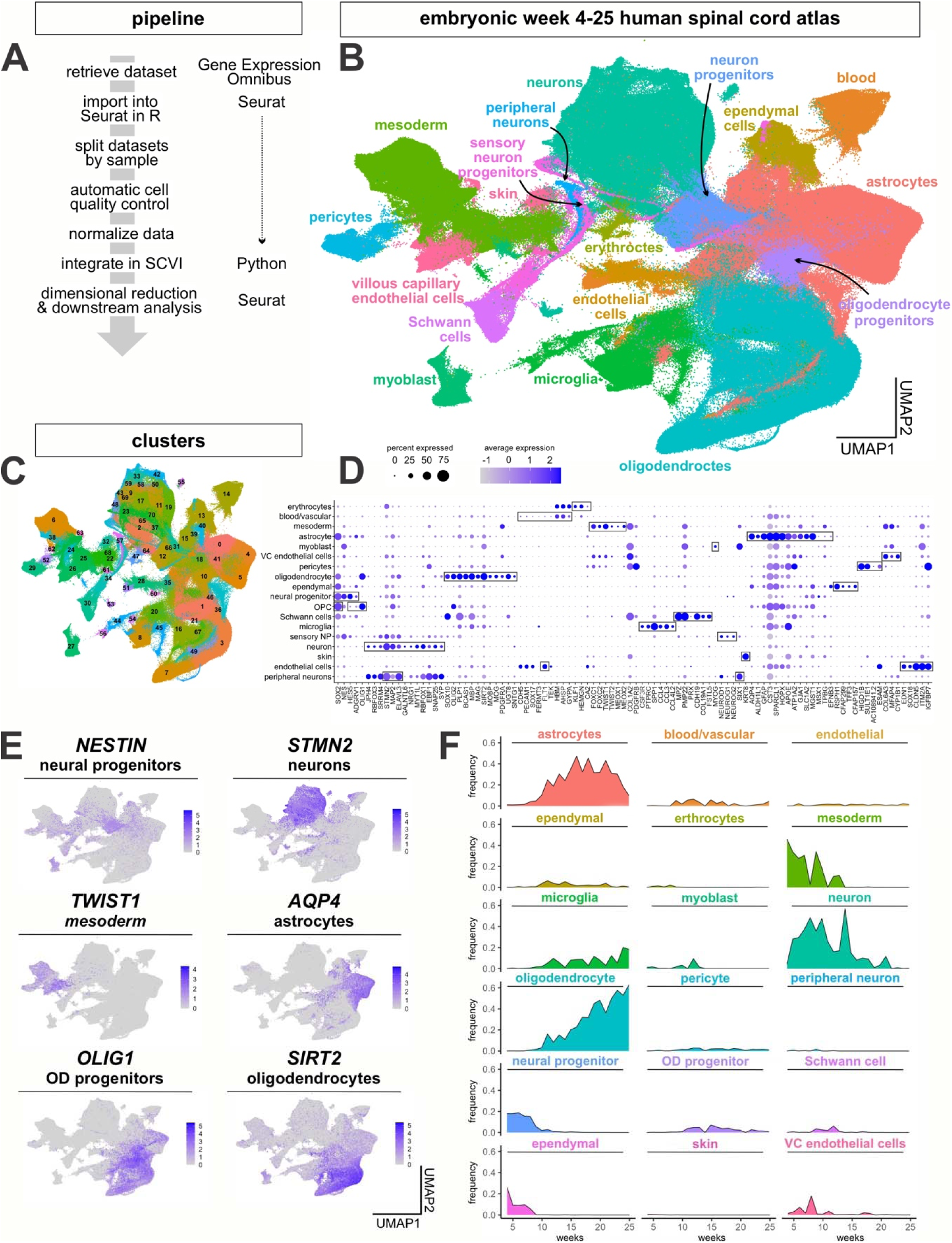
Construction of an *in vivo* human embryonic spinal cord scRNASeq reference atlas. (A) Overview of the analysis pipeline for the single-cell transcriptomic data. (B) The *in vivo* human embryonic spinal cord single cell atlas. Integration of 1.7 million cells from 192 samples spanning gestational weeks 4 to 25 reveals 18 broad categories of cell types present in the spinal cord and surrounding tissues. (C, D) Unsupervised clustering yielded 71 distinct transcriptional clusters (C) that could be grouped into 18 cell types (B), based on their expression of canonical marker genes (boxed genes, D). (E) UMAPs depicting the expression of representative marker genes in the *in vivo* human embryonic spinal cord single-cell atlas. (F) Comparison of the frequency by which the 18 cell types emerge over the timeline of the atlas demonstrates the early proliferation of mesodermal and neuronal cell types, with the later proliferation of astrocytes and oligodendrocytes. The apparent increase in neurons at week 14 likely reflects that all W14 datasets consist of single nuclei samples, whereas datasets from other time points are predominantly single-cell samples.

Clustering analysis using the sctype package^46^ identified 71 transcriptionally distinct populations that were assigned to 18 major spinal cord cell types using curated marker gene sets and combinatorial classification approaches (Fig. 1c,d). These cell classes included neural progenitors, neurons, glia, and mesoderm-derived populations. Canonical lineage markers segregated expected developmental populations with minimal overlap between unrelated cell types (Fig. 1e) and the relative proportion of major cell classes across developmental time recapitulated the expected ontogenetic progression^47, 48^, including early enrichment of progenitor populations followed by sequential emergence of neuronal, astroglial, and oligodendroglial lineages (Fig. 1f). The resulting atlas provides a reference framework for examining lineage diversification and regional specification within the developing human spinal cord.

### The neuronal atlas reveals the expansion and diversification of dI4 and dI5 sensory interneuron populations

To investigate neuronal diversification, we isolated spinal neuronal lineages and reconstructed lineage relationships across developmental time. Integration and reclustering of the neuronal cells generated a high-resolution neuronal atlas containing 308,947 cells, including 56,674 progenitors (Fig. 2a). Using AUCell-based module scoring^49^ and curated gene sets (Fig. 2b, Extended Data Tables 1-2), we assigned all cells dorsal or ventral neuronal identities corresponding to the canonical spinal neuron classes: dorsal interneurons (dI1-dI6), ventral interneurons (v0-v3), and MNs (Fig. 2a).

**Figure 2:**
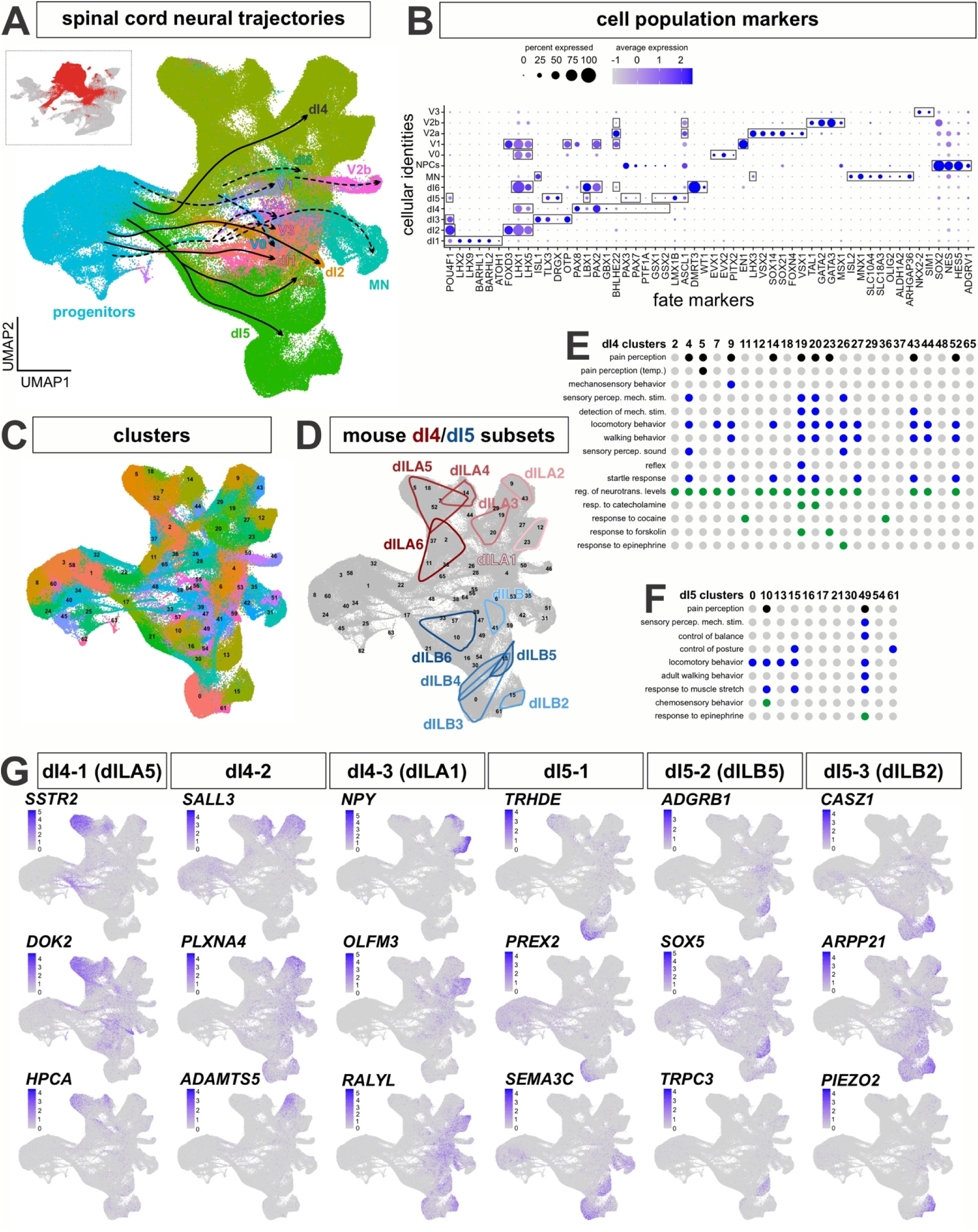
Analysis of dI lineages within the *in vivo* human reference atlas. (A) ∼300,000 neuronal lineage cells were extracted from the overall atlas (inset) and then plotted as 2d UMAP reduction. Labeling cells taken from across the complete timeline with the AUCell classification system identified all the dorsal (dI1-dI6) and ventral (v0-v3, MNs) neural identities in the spinal cord and revealed a vast expansion of the dI4 and dI5 lineages. The differentiation trajectories were hand annotated using gene expression patterns. (B) Dotplot of genes used for classification of each cell type shows expression restricted to expected cell types. (C) Clustering of the neuronal atlas yielded 66 clusters. (D) The expansion of human dI4/dI5 in comparison was assessed in comparison to an embryonic mouse spinal cord single cell atlas^50^. In the mouse study, dI4/dI5 subtypes were designed as late born dILs, dILA1-6 (Extended Data Figure 4A) and dILB1-6 (Extended Data Figure 4B). (E, F) Gene Ontology (GO) enrichment analysis was performed on significantly upregulated genes (avg_log2FC > 0.25) for all clusters identified as either dI4 or dI5 subtypes. Dot plots of GO Biological Process terms found across dI4 (E) and dI5 (F) clusters include various behavioral functions, largely aligning with the canonical roles dI4 and dI5s play in processing sensory information. Black dots identify GO terms related to the sensory perception of pain and temperature; blue dots signify GO terms from different sensory modalities including touch, sound, movement and balance; green dots represent GO terms associated with a response to chemical stimulants. (G) Unique gene markers were identified late-stage subsets of dI4 (dI4-1, dI4-2 and dI4-3) and dI5 (dI5-1, dI5-2 and dI5-3) to further demarcate additional distinct cellular subtypes within the dI4 and dI5 lineages. These genes were selected for their specificity to the cluster, with low to no expression in other clusters.

Trajectory reconstruction identified twelve major spinal neuronal lineages, each characterized by distinct transcriptional programs and lineage relationships (Fig. 2a-c; Extended Data Fig. 3a). Within the dorsal spinal cord, dI4 and dI5 populations emerged as the predominant neuronal classes. Quantitative analysis revealed that dl4 neurons comprised approximately 40% of neurons (∼50% excluding progenitors), while dl5 neurons accounted for approximately 22% (∼30% excluding progenitors) of the neuronal compartment (Fig. 2a). These populations arise alongside other dorsal lineages at approximately gestational week 5 but undergo a pronounced expansion between weeks 7-11 (Extended Data Fig.2), suggesting preferential expansion of these interneuron populations during human spinal cord development.

Subclustering analysis revealed substantial heterogeneity within the dI4/dI5 population, identifying 33 transcriptionally distinct clusters distributed along continuous developmental trajectories (Fig. 2c). Comparative analysis with a recently described mouse embryonic spinal cord atlas^50^ revealed alignment between human dI4/dI5 populations and mouse late-born dILA/dILB populations^51^, supporting the conservation of developmental logic across species (Fig. 2d). Of the 33 human dI4/dI5 clusters, approximately 24 corresponded to mouse counterparts, while nine clusters lacked clear mouse counterparts, suggesting the presence of subclasses unique to the human spinal cord. These findings indicate that the developing human spinal cord exhibits both conserved and species-specific diversification of dorsal sensory interneurons.

### dI4 and dI5 subclasses exhibit transcriptional signatures associated with sensory modalities

We next examined whether the transcriptional heterogeneity within dI4/dI5 populations corresponded to predicted functional specialization. Differential gene expression analysis across dI4/dI5 subclusters followed by gene ontology (GO) enrichment revealed molecular programs associated with multiple sensory modalities, including nociception, mechanosensation, temperature perception, proprioception, and neuromodulatory responses (Fig 2e, f).

Gene ontology analysis further revealed both shared and subtype-specific sensory programs within dI4 and dI5 populations. Shared functions included pain perception, detection of mechanical stimulus (i.e. touch), and regulation of movement, consistent with established roles of dorsal interneurons in somatosensory processing. Several dI4 subclasses exhibited transcriptional signatures associated with temperature sensation (cluster 5), startle response and sensory perception of sound (cluster 4 and 26), suggesting diversification of modality-specific sensory interneuron programs during human spinal cord development. Notably, two dI4 subtypes (clusters 11 and 36) exhibited enrichment in gene programs associated with response to cocaine, supporting our previous observation that embryonic dI4/dI5s may be responsive to psychoactive compounds^32^. These findings suggest that molecular programs associated with neuromodulatory responsiveness are established early during spinal cord development and may represent intrinsic properties of emerging sensory-processing interneuron circuits.

To further resolve functional specialization among dI4/dI5 subclasses, we curated marker gene sets associated with neurotransmission and sensory signaling (Fig. 2i). Several clusters expressed genes with established roles in sensory circuit function. For example, subsets of dl4 populations expressed *SSTR2* (cluster 5, overlapping with mouse dILA5), which modulates neurotransmission^52^ and itch signaling^53^, while *NPY*-expressing populations (cluster 12, overlapping with mouse dILA1/2), encode a peptide neurotransmitter known to inhibit pain perception^54^. Additional subclasses expressed the mechanosensitive ion channel PIEZO2 (cluster 15, overlapping with mouse dILB2) which mediates light touch, proprioception, and mechanical pain^55^. Taken together, these transcriptional signatures indicate progressive specialization of dI4/dI5 interneuron subclasses, including populations associated with neuromodulatory and pharmacologically responsive signaling pathways.

### Human neuromesodermal progenitors (NMPs) exhibit temporally regulated acquisition of posterior axial identity

Given our previous studies with mouse ESCs^32^, we next investigated whether human pluripotent stem cells could generate dIs spanning major axial levels of the spinal cord through an NMP intermediate. Thus, we adapted our NMP-based protocol for human ESCs and additionally examined how extended culture duration and GDF11 treatment^41, 42, 56^ influence dI patterning along the anterior-posterior and dorsal-ventral axes of the spinal cord.

Human NMPs were generated by exposing the H9 ESC line to bFGF and the Wnt agonist CHIR^32, 57^ (Fig. 3a). To examine whether temporal progression influences axial identity *in vitro*, hESCs were exposed to bFGF/CHIR for 2, 4, or 10 days to generate NMP populations with increasingly extended developmental competence (Fig. 3a, b). Immunocytochemical analysis demonstrated expression of canonical NMP markers SOX2, T, and CDX2 across all timepoints (Fig. 3b, c), consistent with bipotential neural–mesenchymal identity.

**Figure 3:**
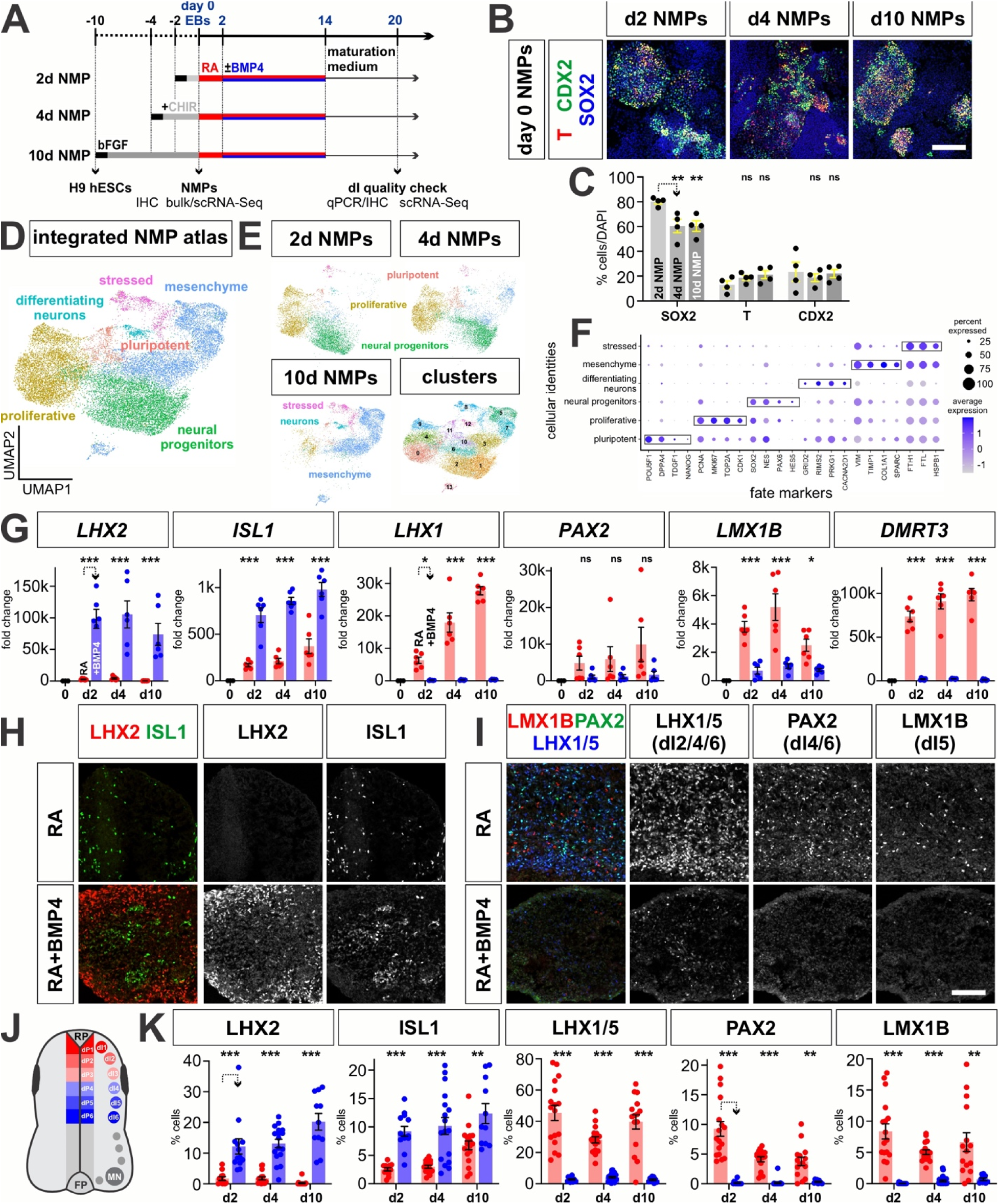
Specification of dI identity in the human RA±BMP4 NMP protocol. (A) Overview of the experimental timeline/workflow for the RA±BMP4 NMP protocols. H9 hESCs were cultured with bFGF/CHIR to induce day (d) 2, 4 and 10 NMPs under 2-dimensional (2D) culture conditions. At day 0, NMPs were sent for both bulk and single cell (sc) RNA sequencing (RNA-Seq) for transcriptional analyses. The NMPs were then converted into 3-dimensional (3D) embryoid bodies (day 0 EBs) and then treated with a 2-day pulse of RA, followed by RA+BMP4 for 12 days to induce dorsal patterning. qPCR and immunohistochemical (IHC) quality checks were performed on day 20. (B, C) IHC analysis of d2 d4 and d10 day 0 NMPs with antibodies directed against T (BRACHYURY, red), CDX2 (green) and SOX2 (blue). Quantification suggests that the number of NMPs is equivalent across the three conditions. (D) UMAP projection of the complete complement of day 0 NMPs, i.e. 2d, 4d and 10d, demonstrates that the cells segregate into neural progenitor and mesenchymal progenitor populations. (E) UMAPs split by culture duration prior to NMP induction depict a progressive shift from neural progenitor enrichment in 2d and 4d NMPs towards mesenchymal identity in 10d NMPs. (F) Dot plot of different NMP fates showing the expression levels of marker genes. (G-I, K) Both qPCR (G) and IHC (H, I, K) analyses of canonical dI markers in day 20 EBs, demonstrate that the RA and RA+BMP4 protocols, using d2, d4 and d10 NMPs, generate the predicted complement of dIs, i.e., RA➔dI4, dI5, dI6 and RA+BMP4 ➔ dI1 and dI3. (J) Schematic representation of the developing spinal cord *in vivo*, depicting the location of the six populations of dorsal progenitors (dP) and dorsal interneurons (dIs). Scale bar: 100µm (B); 75µm (H, I) Probability of similarity between control and experimental groups: *= p < 0.05, **p<0.005 *** p<0.0005; two-way ANOVA.

Single-cell and bulk RNA sequencing revealed greater transcriptional divergence from pluripotent states as NMP culture duration increased (Fig. 3d-f, Fig. 4e-o). GO analyses indicated gradual enrichment of mesoderm-associated transcriptional programs and reduction of anterior neural signatures with extended culture duration (Fig. 4g-n). Consistent with posteriorization over time, HOX gene expression increased with longer NMP culture duration, with 2-day NMPs expressing anterior HOX genes such as HOXC5, while 4- and 10-day NMPs expressed progressively more posterior HOX paralogs, including HOXA10, HOXD11, and HOXD13 (Fig. 4o). These findings indicate that human NMPs can acquire posterior competence through temporal progression in culture.

**Figure 4:**
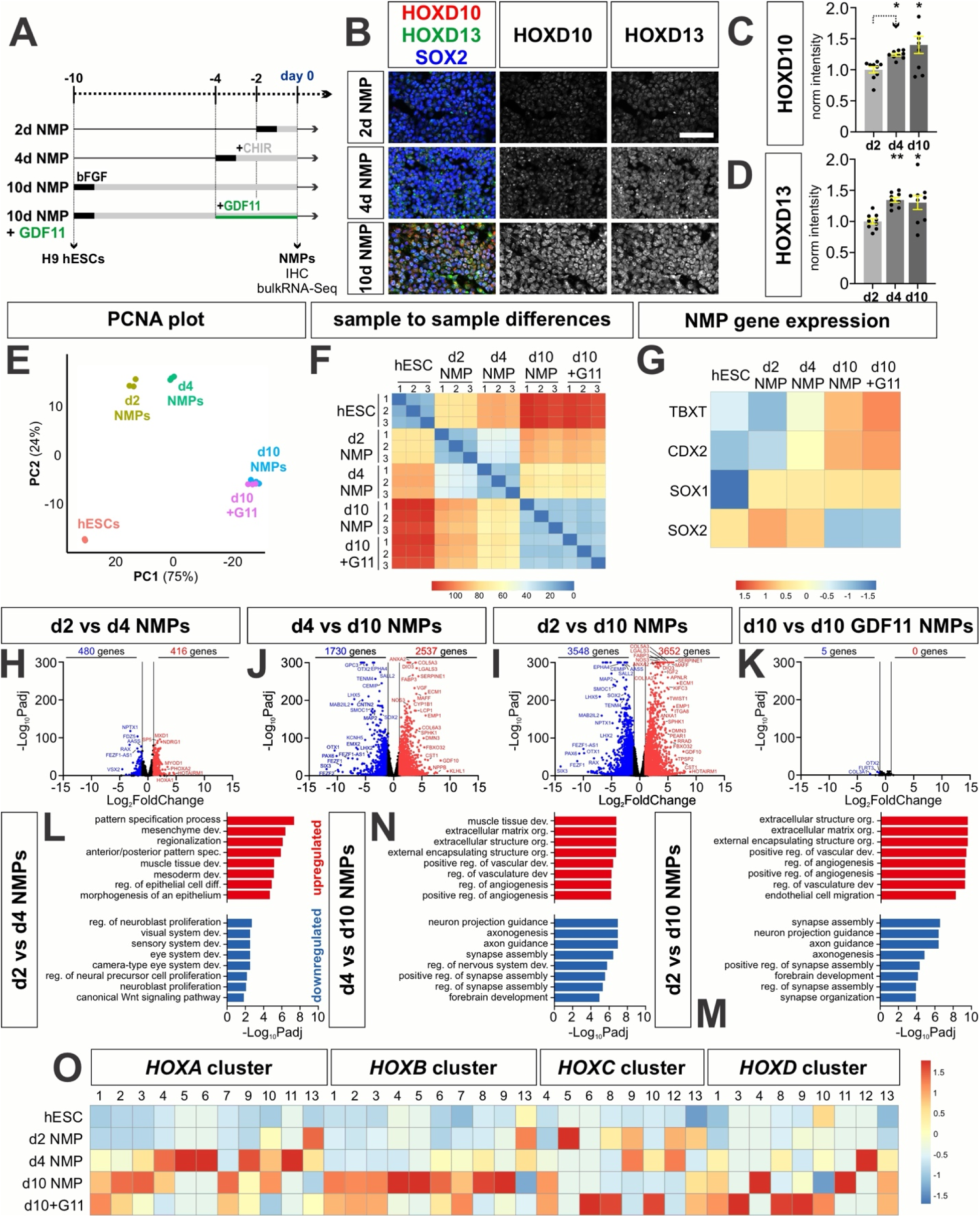
Assessing the effect of culture duration on NMP identity along the anterior-posterior axis. (A) Overview of the experimental timeline/workflow for the RA±BMP4 NMP protocols. (B -D) IHC analyses of NMPs immediately before EB formation also suggests that prolonging the culture time progressively posteriorizes NMPs. Two posterior HOX genes - HOXD10, and HOXD13 - have significantly higher expression in d10 NMPs, compared to d2 NMPs. SOX2 levels are ∼constant across the d2, d4 and d10 NMP conditions, indicating that the NMP identity is maintained. (E) Principal-component analysis (PCA) of bulk RNA-seq expression profiles from hESCs, d2, d4, d10, and d10+GDF11 NMPs shows that samples segregate along PC1 (75%) according to the duration of NMP induction, suggesting culture time is the primary driver of transcriptional variation. (F) Heatmap of pairwise Euclidean distances between samples. hESCs are most dissimilar from d10 and d10 + GDF11 NMPs, while d10 and d10+GDF11 samples are highly transcriptionally similar. (H-K) Volcano plots comparing the differentially expressed genes (DEGs) in the d2, d4 and d10 NMP cultures (|log□ fold change| ≥ 1, Benjamini-Hochberg-adjusted p < 0.05). The magnitude and number of transcriptional differences increase with greater separation in induction time, with the largest number of DEGs observed between d2 and d10 NMPs, and minimal differences between d10 and d10+GDF11 NMPs, indicating largely shared transcriptional states. (L-M) Gene Ontology (GO) enrichment analysis of significantly upregulated and downregulated genes in contrasts. Genes upregulated in d4 NMPs are enriched for patterning and morphogenetic processes, while d10 NMPs show strong enrichment for extracellular matrix organization and vasculature-associated pathways. Downregulated gene sets suggest that forebrain-associated transcriptional programs are progressive reduced with increased culture time. (O) Heatmap of average *HOX* gene expression across conditions, scaled per gene. Robust *HOX* expression emerges in d4 NMPs, predominantly in the HOXA cluster. d10 NMPs show strong *HOXB* and *HOXD* gene expression, while the addition of GDF11 increases expression in the *HOXC* and *HOXD* clusters, suggesting that prolonged culture duration and GDF11 exposure play a role modulating axial identity through differential activation of *HOX* clusters. Scale bar: 20µm Probability of similarity: *= p < 0.05, **p<0.005; two-way ANOVA.

### Human NMP-derived cultures generate the full complement of dI identities

We next examined whether temporally patterned NMPs could generate the full range of dI subtypes. NMPs were aggregated into embryoid bodies (EBs) and neuralized with retinoic acid (RA) to promote spinal neural identity (Fig. 3a). qPCR and IHC analyses demonstrated robust generation of intermediate dI populations (dI4-dI6) across all NMP timepoints (Fig. 3g-k, Extended Data Fig. 4e). Addition of BMP4 during neuralization promoted specification of dorsal-most interneuron populations (dI1-dI3), consistent with the known developmental signaling logic^32^. Optimization of BMP4 timing defined day 2–14 as an effective window for dorsal specification (Fig. 3a, Extended Data Fig. 4a-d). Combined RA+BMP4 treatment generated dI1 and dI3 populations across NMP conditions (Fig. 3g-k, Extended Data Fig. 4e), while scRNA-seq analyses further identified dI2s not readily detected using qPCR or IHC (Fig. 5g; Fig. 6a). Collectively, these results demonstrate that temporally patterned human NMP cultures generate the full complement of dI classes spanning multiple axial identities and reproduce key features of *in vivo* HOX-dependent spinal cord patterning.

**Figure 5:**
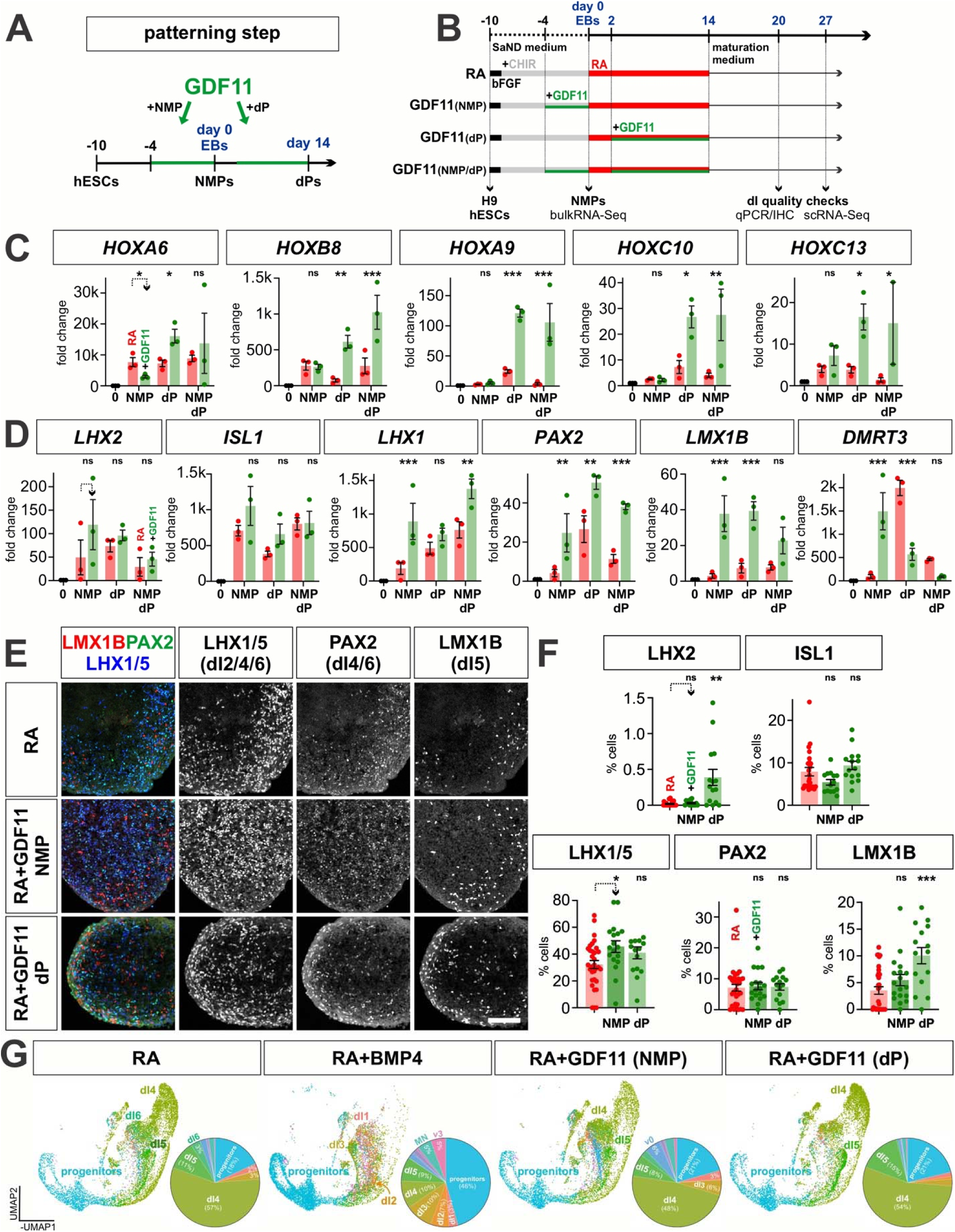
Assessing the effect of GDF11 treatment on both dorsal-ventral and anterior-posterior identity. (A, B) Overview of the experimental timeline/workflow for the RA±GDF11 NMP protocols. The effect of GDF11 treatment was assessed on NMP patterning, dP patterning and both NMP and dP patterning (A). (C) qPCR analyses of a range of *HOX* gene expression suggests that GDF11 addition at the dP stage is the most effective at driving the expression of the posterior most HOX genes (*HOXA9*, *HOXC10* and *HOXC13)*. (D-F) qPCR analyses of day 20 EBs (D), suggest that the addition of GDF11 at either the NMP or dP stage significantly increases expression of *PAX2* and *LMX1B*, the canonical genes that mark the dI4/dI5 identities. However, the IHC analysis suggest the effect of GDF7 on protein levels is more subtle, with the most significant effect observed when GDF11 added at the dP stage on dI5 identities. (G) UMAPs of day 27 *in vitr*o-derived cells generated using the RA, RA+BMP4, and RA+GDF11 (NMP and dP) protocols, using neural spinal cord cell-type identity transferred from the *in vivo* neuronal atlas (Fig. 2a). The pie charts summarize the proportions of neuronal cells generated by each protocol. The RA and RA+GDF11 conditions predominantly yield dI4-dI6 identities, whereas the addition of BMP4 shifts patterning towards the dorsalmost dI1-dI3 identities. Adding GDF11 during NMP induction has minimal effects on patterning compared to RA alone, while GDF11 treatment during dP formation increases in number of dI5s, apparently at the expense of the dI6 population. Scale bar: 75µm Probability of similarity between control and experimental groups: *= p < 0.05, **p<0.005 *** p<0.0005; two-way ANOVA

**Figure 6:**
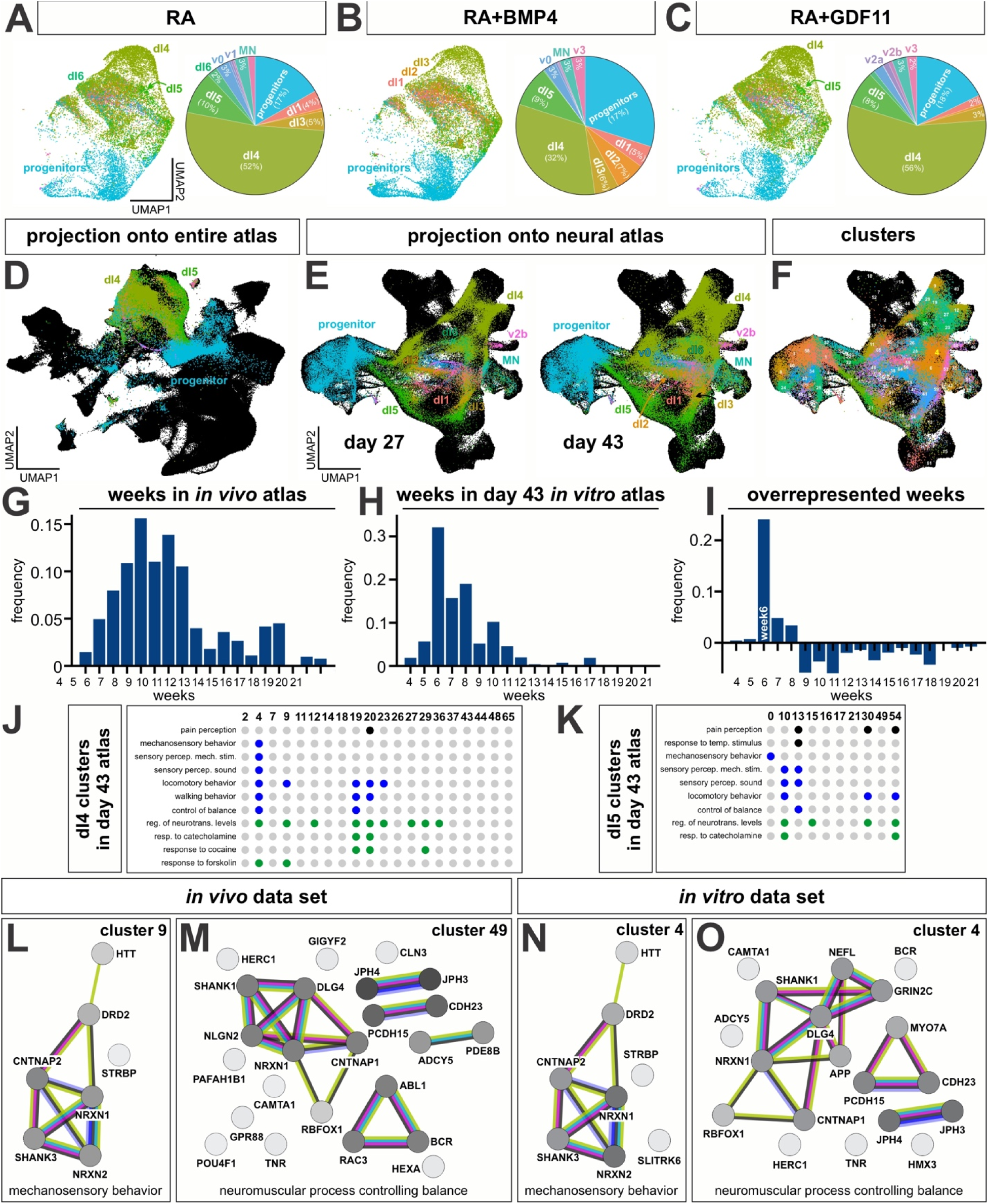
Comparison of *in vitro* data with the *in vivo* reference atlas reveals ASD signatures. (A-C) UMAPs of day 43 *in vitr*o-derived cells generated using the RA, RA+BMP4, and RA+GDF11 (NMP) protocols, using neural spinal cord cell-type identity transferred from the *in vivo* neuronal atlas (Fig. 2a). The pie charts summarize the proportions of neuronal cells generated by each protocol. The RA condition predominantly yields dI4-dI6 identities, while the RA+BMP4 protocol shifts patterning toward dI1-dI3 populations. Addition of GDF11 prior to NMP induction does not alter resultant cell-type composition compared to RA alone. (D) Projection of day 43 *in vitro* cells onto the complete *in vivo* spinal cord reference (shown as a black silhouette). I*n vitro* neural progenitors and neurons map directly onto their *in vivo* counterparts, selectively recapitulating endogenous spinal neural transcriptional identities. (E, F) Projection of day 27 and 43 *in vitro-*derived cells onto the neuronal subset of the *in vivo* spinal cord atlas. The *in vitro* dIs map along the correct lineage trajectories (E) suggesting they are transcriptionally indistinguishable from endogenous neurons. The *in vitro* dIs are colored by transferred neuronal cell-type identity (E) or atlas cluster identity (F). (G-I) The day 43 *in vitro* datasets (G) align most closely with week 6 samples in the *in vivo* reference atlas (H). Using the relative frequency of the developmental weeks in the *in vivo* atlas as a reference, we assessed for overrepresentation and found that week 6 profiles were enriched in the *in vitro* dataset compared with the *in vivo* atlas (I). (J, K) Gene Ontology (GO) enrichment analysis was performed on significantly upregulated genes (avg_log2FC > 0.25) for any stem cell derived clusters assigned to a dI4 or dI5 atlas cluster, as performed previously for the *in vivo* dI4/dI5 clusters. Dot plots of enriched GO Biological Process terms in the dI4 (J) and dI5 (K) clusters include behavioral categories that overlap with those observed in the in vivo datasets (Fig. 2e, f). Black dots are related to the sensory perception of pain and temperature, blue dots correspond to sensory modalities including touch, sound, movement and balance, while green dots represent a response to chemical stimulants. (L-O) To assess the extent to which *in vitro* and *in vivo* dIs may be functionally similar, the genes making up each GO term were subjected to a STRING analysis, thereby identifying the protein interaction networks that encode specific sensory behavioral modalities. These networks include a mechanosensory circuit derived from *in vivo* cluster 9 (L), as well as a balance regulating circuit derived from *in vivo* cluster 49 (M), include proteins with variants that was strongly associated with autism spectrum disorder: NLGN2, NRXN1, SHANK1/3, DLG4 and CNTNAP2. These circuits were recapitulated in the *in vitro* cluster 4 mechanosensory behavior protein interaction network (N) and the *in vitro* cluster 4 balance regulating protein interaction network (O). For STRING networks (L-O), nodes represent proteins, edges represent protein-protein associations. Edge color indicates known interactions; experimentally determined (purple), from curated databases (teal), predicted interactions; gene neighborhood (green), gene fusions (red), gene co-occurrence (blue) and others; textmining (yellow), co-expression (black), protein homology (periwinkle)

### GDF11 reinforces posterior identity and selectively influences dI specification

We next investigated whether GDF11, a regulator of posterior HOX gene expression in the neural tube^43^, contributes to axial patterning or dI specification in human NMP-derived cultures^41, 42, 56^. GDF11 was applied either during the NMP stage, during dorsal progenitor (dP) differentiation, or at both developmental stages (Fig. 5a, b). Bulk RNA-Seq and qPCR analysis revealed that GDF11 applied during the NMP stage had minimal impact on global transcriptional identity (Fig. 4e,k), although subtle differences in *HOX* gene expression profiles were observed (Fig. 4o). In contrast, GDF11 applied during dorsal progenitor differentiation increased the expression of posterior HOX genes including *HOXA9*, *HOXC10* and *HOXA13*, consistent with reinforcing lumbosacral identity (Fig. 5c).

GDF11 treatment also modulates dI specification, in a temporally restricted manner. While GDF11 did not significantly alter generation of dorsal-most interneurons, dI1–dI3, it increased expression of markers associated with the dl4 and dl5 populations, compared to the RA condition (Fig. 5d-f; Extended Data Fig. 7k). GDF11 treatment at the dorsal progenitor, but not NMP, stage increased the number of LMX1B^+^ dI5s in both IHC and scRNA-Seq analyses (Fig. 5e-g). GDF11-dP can also drive the formation of more PAX2^+^ dI4s (Extended Data Fig. 7k), although this result is inconsistent across replicates (Fig. 5f). GDF11-dP also suppresses *DMRT3* expression, the canonical marker for dI6s, suggesting reduced numbers of dI6s population (Fig. 5d, g). For all conditions, there is no synergistic effect of adding GDF11 at both the NMP and dP stages, and GDF11 generally acts to suppress the effect of BMP4 addition (Extended Data Fig. 5). Taken together, these findings suggest that GDF11 functions as a stage-dependent modulatory signal that reinforces posterior identity and promotes the selective specification of dI subtypes.

### Stem cell-derived dIs align with endogenous developmental trajectories

To determine whether stem cell-derived dIs recapitulate endogenous developmental programs, neurons generated under RA, RA+BMP4 and RA+GDF11 conditions were analyzed by scRNA-Seq at day 43 (Fig. 6a-c). All differentiation conditions generated substantial populations of dI4 (50%) and dI5s (∼10%), consistent with the lineage distribution observed in the *in vivo* atlas. RA+ BMP4 treatment increased representation of dorsal-most interneurons, including dI2 populations, while small populations of ventral interneurons and MNs were also detected, suggesting emergence of intrinsic patterning centers within the EBs .

Using scVI and scArches^58^, we projected the *in vitro* single-cell datasets onto the full *in vivo* reference atlas (Fig. 1b) and the *in vivo* neuronal-specific atlas (Fig. 2a, Fig. 6d, e). This approach revealed strong alignment of stem cell-derived neurons with endogenous spinal neuron trajectories (Fig. 6d, e). Label transfer analysis demonstrated the preferential alignment of *in vitro*-derived cells with neuronal populations rather than non-neuronal lineages, supporting faithful recapitulation of spinal neuronal identities. Specifically, the day 27 and day 43 *in vitro*-derived dIs map along the corresponding *in vivo* lineage trajectories (Fig. 2a, 6e). Consistent with day 43 dataset being slightly more mature, the number of dI4/dI5s are increased compared to the day 27 dataset and they extend further along their trajectories (Fig. 6e). Taken together, this data suggests that the stem-cell derived dIs are transcriptionally indistinguishable from their endogenous counterparts.

While the *in vitro* dataset overlaps primarily with regions present in the earliest time points in the atlas (i.e., weeks 4-7), it is underrepresented in the dI4/dI5s specified from week 8 and onward (Fig. 6f, Extended Data Fig. 2). To more formally assess the maturation state of the *in vitro* neurons, we trained a classifier on the week metadata in the scVI space of the neural atlas and transferred these labels to our *in vitro* dataset. Comparison of developmental stage signatures indicated that day 43 *in vitro* neurons most closely resembled gestational week 6 populations (Fig. 6g-i), consistent with expected maturation timelines. These findings indicate that human stem cell-derived dIs align with endogenous lineage trajectories and developmental stage.

### Mechanosensory interneuron networks enriched for ASD-associated genes emerge during early spinal cord development

We next investigated whether transcriptional programs associated with dI4/5 subclasses provide insight into the organization of the developing sensory circuits. Projection of datasets from *in vitro-*derived neurons onto the neural atlas enabled assignment of cells to defined dI4/dI5 clusters, facilitating comparison of predicted functional programs across populations (Fig. 1c, 6f).

GO analyses identified conserved molecular programs associated with pain perception, mechanosensation, neurotransmitter regulation, and balance across *in vitro* and *in vivo* datasets (Fig. 6j,k). Protein interaction network analysis using STRING^59^ identified conserved molecular modules that encode pain perception (dI4s, Extended Fig. 8a, d, e, h, i; dI5s, Extended Fig. 10a, b, c, f), the response to cocaine (dI4s, Extended Fig. 8b, c, f), perception of mechanical stimulus (dI4s, Extended Figs. 8e and 9a, b, d, e; dI5s, Extended Fig. 10e) mechanosensory behavior (dI4s, Fig. 7l, n) and balance (dI4s, Fig. 7o, Extended Fig. 9c; dI5s, Fig. 7m). While day 43 *in vitro* neurons are relatively immature (Fig. 6h, i), several networks share molecular signatures with *in vivo* networks encoding the same function. For example, networks encoding pain perception in cluster 20 share a protein loop centered on NPY1R, thought to mediate neuropathic pain^60^ (Extended Fig. 8d, e), while a dI4 touch circuit in cluster 4 (Extended Fig. 9b) resembles an *in vivo* dI4 touch circuit within the corresponding cluster (Extended Fig. 9a).

Several clusters associated with mechanosensory behaviors (Fig. 6n) and balance (Fig, 6o) exhibited enrichment of genes with high SFARI scores^15^, including NGLN2^61, 62^, NRXN1/2^61, 63^, SHANK1/3^64^, DLG3^65^, CNTNAP2^66^. Identical transcriptional signatures were observed in both *in vivo* and *in vitro* dI4/dI5 populations and emerged at very early developmental stages, i.e., gestational week 6.

Given the frequent occurrence of altered sensory processing and balance phenotypes in individuals with ASD, these findings suggest that molecular programs associated with neurodevelopmental disorders are present within developing spinal cord sensory circuits. These results raise the possibility that sensory phenotypes associated with neurodevelopmental disorders may arise from distributed circuit-level developmental programs spanning both spinal and supraspinal regions.

## Discussion

In this study, we establish multiple human NMP–based directed differentiation strategies that generate dorsal spinal interneurons across axial levels and benchmark their identities against a temporally resolved single-cell atlas of the developing human spinal cord. Integration of *in vitro* and *in vivo* datasets demonstrates that NMP–derived dIs recapitulate endogenous transcriptional identities and developmental trajectories, enabling subtype-level comparison between stem cell-derived and fetal spinal neurons. The atlas further reveals extensive diversification of dI4 and dI5 populations, identifies molecular programs associated with sensory-processing functions, and uncover transcriptional networks enriched for high-confidence ASD risk genes. Together, these findings provide a framework for understanding how human dorsal spinal interneurons acquire positional and functional identity.

### Temporal competence and signaling context jointly shape posterior identity

Our results support a model in which posterior spinal cord identity emerges through the interaction between developmental time and signaling environment rather than via a single instructive cue. Extending NMP culture duration promotes the sequential acquisition of posterior competence, consistent with previous work showing that temporal progression contributes to HOX gene activation^41^. However, we also find that extended time in culture alters NMP identity in more complex ways, such as inducing the progressive enrichment of mesodermal gene signatures (Fig. 4). Thus, while posterior competence increases with time, prolonged NMP maintenance may simultaneously bias lineages, underscoring a tradeoff between axial maturation and progenitor plasticity.

We observe broad and overlapping *HOX* gene expression profiles, consistent with previous work^41, 67^, suggesting that posterior identity in human neural progenitors may reflect combinatorial *HOX* activity rather than sequential activation of posterior *HOX* paralogs (Fig. 4). Similar deviations from classical HOX co-linearity have been observed in analyses of human fetal spinal cord development^18^, supporting the idea that human axial patterning involves more flexible regulatory logic than inferred from model organisms.

Our analysis also reveals context-dependent effects of GDF11. While GDF11 applied during NMP induction has modest effects on transcription, GDF11 applied during dorsal progenitor differentiation reinforces posterior HOX expression and selectively influences dI subtype specification. These findings suggest that GDF11 acts as a modulatory signal that stabilizes lineage decisions once progenitors have acquired sufficient temporal competence. Such temporally gated responsiveness may reflect prolonged developmental plasticity characteristic of human neural progenitors^68^ and highlights how human spinal cord patterning may differ from signaling hierarchies derived from rapidly developing model organisms with shorter gestation periods.

### Expansion of dI4 and dI5 populations reveals diversification of human somatosensory interneurons

The integrated human spinal cord atlas provides a temporally resolved view of interneuron diversification and reveals a striking expansion of dI4 and dI5 populations at later gestational stages (Fig. 2a and Extended Data Fig.- 2). Comparative analysis with mouse embryonic spinal cord atlases^50^ indicates substantial overlap between human dI4/dI5 clusters and late-born mouse dIL_A_/dIL_B_ populations, suggesting conserved developmental origins.

dI4s and dI5s have transcriptional signatures that are associated with molecular pathways implicated in mechanosensation, nociception, and neuromodulatory signaling. For example, subsets of dI5 populations express PIEZO2^68^, a mechanosensitive ion channel required for touch and proprioception, while subsets of dI4 neurons expresses NPY, a neuropeptide associated with inhibitory modulation of pain pathways. We also identify transcriptionally dI4/dI5 subclusters with human specific sensory functions, consistent with observed species-specific differences in somatosensory modalities^69^. Moreover, the mechanistic basis for individual somatosensory differences remains an intriguing and understudied area. Subtle biases within dI4/dI5 populations could potentially explain wide-ranging differences between individuals, such as pain sensitivity or proprioceptive acuity. Such diversification underscores the importance of human developmental atlases for identifying cell types and regulatory programs not completely captured in model systems.

### Developmental origins of sensory phenotypes associated with neurodevelopmental disorders

An unexpected finding of the atlas is the enrichment of ASD-associated gene networks within the dI4/dI5 interneuron populations (Fig. 6-o, Extended Data Figs. 8-10). Both *in vitro* and *in vivo* dI4/dI5 clusters exhibit gene networks associated with mechanosensation, balance, and neuromodulatory signaling that include genes strongly linked to ASD risk, including NRXN1^70^, SHANK1/3^71^, DLG4^72^, CNTNAP2^73^, and NLGN2^74^. Although ASD is primarily studied in the context of cortical development, sensory processing differences are a prominent feature of the disorder.

The presence of ASD-associated molecular signatures in developing spinal interneurons raises the possibility that sensory phenotypes associated with ASD (such as differences in touch and balance processing) may arise from developmental programs spanning both spinal and supraspinal circuits. Notably, these transcriptional signatures are detectable in stem cell-derived interneurons at early developmental stages, consistent with the idea that ASD-associated molecular differences emerge during embryonic circuit formation ^75^. While further functional studies are required to establish causal relationships, these findings highlight the potential contribution of spinal sensory circuitry to neurodevelopmental phenotypes conventionally attributed primarily to brain-specific mechanisms.

### Implications for regenerative strategies targeting sensory circuits

The ability to generate dI subclasses matched for axial identity and developmental stage has implications for regenerative medicine and SCI research. Sensory dysfunction following SCI reflects both neuronal loss and disruption of dI circuits that integrate mechanosensory information. The reference atlas indicates that dI4 and dI5 populations represent major components of human dorsal sensory circuitry and our differentiation protocols recapitulate this lineage distribution.

The close transcriptional correspondence between stem cell-derived and endogenous dIs suggests that NMP-based differentiation strategies can produce cell populations that approximate physiologically relevant sensory interneuron identities. Such lineage-matched populations provide a rational starting point for reconstructing dorsal spinal circuits disrupted by injury. More broadly, the ability to generate defined interneuron subclasses provides a foundation for mechanistic studies of human sensory circuit assembly and for testing cell replacement strategies designed to restore sensory function.

### Limitations and future directions

Several questions remain regarding the specification and maturation of human dIs. Some populations, including dI2s, remain difficult to detect using conventional marker-based approaches and may represent transient developmental states or require additional patterning cues. In addition, the *HOX* gene expression patterns *in vitro* do not follow a simple anterior–posterior hierarchy, suggesting that axial identity in human neural progenitors may be governed by combinatorial or context-dependent regulatory mechanisms. Further studies will be required to determine how maturation state influences the functional integration of stem cell-derived interneurons and whether these cells can restore circuit function in models of spinal cord injury. Together, our findings highlight principles governing specification of human dorsal spinal interneurons and establish principles for engineering sensory circuit cell types with defined positional and functional identities.

## Methods

### Construction of the human atlas

#### Integration of single-cell datasets from different origins

Data from six publications^18, 36–40^ were loaded into Seurat v5^44^ in R. For each dataset, all samples were combined into one object, and each sample was split into a separate layer. Quality control was performed automatically by removing cells for each sample that fell outside of three median absolute deviations (both sides for nCount_RNA and nFeature_RNA, and higher for percent.mt). The Anderson dataset did not include mitochondrial genes and thus was only filtered on the two other metrics. After QC, all datasets were combined into one large Seurat object and the data was normalized and variable features were calculated. The object was then converted to anndata and integration was performed using SCVI^45^ on the counts layer using only the variable features (n_layers = as.integer(2), n_latent = as.integer(30), gene_likelihood = "nb"). The latent representation was then brought back into the full Seurat object and clusters and UMAP were calculated. Cluster identities were assigned using sctype^46^ and a custom list of marker genes collated from various publications.

### Reclustering of neuronal clusters

To analyze the neuronal population, clusters identified as neurons, peripheral neurons, sensory neuron progenitors, and progenitors-neurons were isolated from the atlas and reintegrated using newly calculated variable features and new scVI parameters as recommended by scArches^58^ (use_layer_norm = "both", use_batch_norm = "none", encode_covariates = T, n_layers = as.integer(2), dropout_rate = as.numeric(0.2)). After dimensional reduction, a few undesired populations were still left, so clusters that were identified either as peripheral populations, or unidentified neuronal populations (mostly identified by a lack of distinct marker genes) were removed and the dataset was reintegrated.

### dI cell type extraction based on the marker gene expression

To identify the various populations of cells within the neuronal atlas, AUCell^49^ was used with a custom list of marker genes to create a module score for each cell, for each possible cell type. Cutoffs were manually assigned for scores for each cell type, and every cell was assigned 0, 1, or multiple identities. Cells with multiple identities were then rectified using a kNN classifier (k=10) to change their identity to match those of surrounding cells. Then the same was done with cells assigned no identity to extrapolate to nearby populations not expressing high levels of any module.

### dI4/dI5 subcluster analysis

Differential gene expression analysis across clusters was performed using the FindAllMarkers function from the Seurat R toolkit. The results were filtered, to only include clusters identified as containing large portions of either dI4 (2, 4, 5, 7, 9, 11, 12, 14, 18, 19, 20, 23, 26, 27, 29, 36, 37, 43, 44, 48, 52, 65) or dI5 (0, 10, 13, 15, 16, 17, 21, 30, 49, 54, 61). Of note is that due to the AUCell classification process, clusters are not made up exclusively of one cell type, but each cluster identified here has a large portion of cells identified as either dI4/5. Gene Ontology (GO) enrichment analysis was performed on significantly upregulated genes in each cluster (avg_log2FC > 0.25) using the enrichGO function from the clusterProfiler R package. The ontology category was restricted to Biological Process (BP), multiple testing correction was applied using the Benjamini-Hochberg method, and terms were considered significant with an adjusted p-value of 0.05 and q-value of 0.2. GO categories for dot plot visualization were selected based on their association with various sensory modalities and overlap, or lack thereof, amongst clusters. The STRING (Search Tool for the Retrieval of Interacting Genes/Proteins) resource was utilized to explore known and predicted protein-protein interaction networks from genes that were making up GO categories of interest.

### Single-cell RNA sequencing and analysis of in vitro cultures

#### Preparation of single cell suspension

Single cell suspension from EBs is generated via papain dissociation method using the Worthington papain dissociation kit (refer to <u>manual</u>). Briefly, EBs are first collected in 15mL centrifuge tubes and washed twice with DPBS. The papain solution is prepared by mixing papain with DNAse (supplied in the kit), and the solution is allowed to activate for 10 minutes at 37°C before adding to the EBs followed by triturating 10 times with a p1000 pipette. EBs are then preferably placed on an orbital shaker at 37°C and 5% CO_2_ for 1 hour and 15 minutes and triturated 10 times every 5 minutes or until no visible EB clumps remain. Single-cell suspension of EBs are then centrifuged at 300 x g for 7 minutes. The cell pellet is resuspended in 2mLs of Ovomucoid protease inhibitor solution to stop the papain digestion and passed through a 40µm cell strainer to remove undigested organoid debris. Single cells are then centrifuged once more at 300 x g for 7 minutes and the cell pellet is then resuspended in 1mL of 0.04% BSA in DPBS for GEM preparation using 10X genomics platform (the concentration of BSA can be increased to reduce cell stickiness/chance of doublets).

### Library preparation, and sequencing

∼ 10,000 live cells/condition were used to generate single-cell cDNA libraries using protocol described by the 10X Genomics 3’GEX library preparation kit. Cells were first partitioned into GEMs (gel-beads-in-emulsion) containing using the 10x Genomics Chromium controller. Further, 3’ mRNA-seq gene libraries were prepared using the Chromium Single Cell 3′ Library & Gel Bead Kit v2 (10x Genomics), according to the manufacturer’s instructions. Library quality control was assessed using Agilent and Qubit assays. If quality control is within acceptable conditions (e.g., library concentration of 20-40ng/ul), libraries were sequenced on the Illumina NovaSeq X Plus 10B. For 3’GEX libraries targeting 10,000 cells 1,800 million reads/sample were obtained. The GEO accession numbers for all the datasets can be found in Extended Data Table 3.

### Integration of single-cell in vitro datasets

Single-cell libraries were processed with the 10x Genomics Cell Ranger Count pipeline (v9.0.1) and loaded into Seurat v5 in R. Within the Seurat object, quality control was performed for each sample by removing cells that fell outside of three median absolute deviations for total UMI counts (nCount_RNA) and number of detected genes (nFeature_RNA), and above three median absolute deviations for percent mitochondrial reads (percent.mt). After QC, data were normalized and variable features calculated.

For day 43 samples, separate Seurat objects were created for the RA/BMP series without GDF11 and the RA ± GDF11 experimental series. Following QC, data were normalized and variable features calculated independently before the two objects were merged into a single Seurat object for subsequent integration.

For nonlinear batch integration, the Seurat object was subset to variable features and converted to an AnnData object. Integration was performed with SCVI, using a model trained on raw counts with default settings (batch_key = "sample"), to obtain a low-dimensional latent embedding. The latent representation was imported back into Seurat as a dimensional reduction, and was used to compute nearest-neighbor graphs, clusters, and UMAP embeddings. To provide an independent annotation of broad neuronal versus non-neuronal populations in the in vitro cultures, we applied sctype with a custom list of marker genes collated from various publications. For downstream atlas-based analyses, the dataset was restricted to cells classified by sctype as neuron, peripheral neuron, sensory neuron progenitor, or progenitor–neuron. Populations with overwhelmingly stressed marker genes or a lack of distinct marker genes were additionally removed. Examples of scRNA-Seq datasets before the neural lineage restriction are included in Extended Data Fig.7A-J.

### Projection of in vitro cells onto the human atlas

Projection of the in vitro cells onto the human spinal cord atlas and its neuronal subset was performed using pretrained SCVI reference models built on the whole in vivo atlas and the neuronal atlas respectively. The in vitro Seurat object was subset so that its gene set matched the genes present in the reference AnnData object. After conversion of the object to AnnData, reference mapping followed the scArches workflow (SCVI.prepare_query_anndata followed by SCVI.load_query_data), and the query model was fine-tuned for 200 epochs to align the in vitro dataset to the reference latent space. The resulting query-aligned latent representation was imported into Seurat and projected into the appropriate reference UMAP space using the saved UMAP models stored in the atlas objects.

To transfer annotations from the *in vivo* neuronal atlas to the in vitro cells, we trained supervised machine-learning models using atlas SCVI embeddings as predictors. K-nearest neighbors (k = 10) was used for dI cell-type classification (based on AUCell-derived identities; see dI cell type extraction based on the marker gene expression) and prediction of developmental week, while random forest (ranger) was used for cluster identity transfer. Cluster-specific markers for cells assigned to in vivo dI4 or dI5 enriched clusters were identified using FindAllMarkers. These gene sets were used to assess if in vitro cells recapitulate in vivo dI4/dI5 subtype transcriptional programs (see dI4/dI5 subcluster analysis).

### Bulk-RNA sequencing and data processing

#### RNA preparation

NMP cultures subjected to bulk-RNA sequencing were first washed with 1X DPBS to remove the media. Cells were then lysed by directly adding 600μL of RLT buffer in the culture wells, and the lysate is then collected and flash frozen with liquid nitrogen to be stored at -80°C. The total RNA was then purified using Qiagen’s RNeasy kit, and eluted in 50μL of RNAse-free water. Using a microvolume spectrophotometer, RNA concentrations were determined and diluted to 1μg/μL. RNA samples were further subjected to quality control assessment using Agilent Technologies 2100 Bioanalyzer and only samples with a RIN score >8.0 were used for library construction using Universal plus mRNA sequencing kit (NuGEN). Libraries were then sequenced on an Illumina Novaseq X Plus to generate ∼30 million reads per sample. The GEO accession numbers for all the datasets can be found in Extended Data Table 3.

### Processing and differential expression analysis

Reads were obtained as raw FASTQ files, assessed for quality with FASTQC, and aligned to the human reference genome (GRCh38, GENCODE v29 annotation) using STAR. Gene-level counts were obtained with featureCounts (exon-level, paired-end read counting). Differential gene expression was tested using DESeq2. For each reference level of interest (hESC, D2NMP, D4NMP, D10NMP), a re-leveled model was built and processed separately using the design formula ∼ condition, to obtain pairwise contrasts across all conditions. Global transcriptional relationships among samples were assessed using VST-normalized counts from the hESC-referenced model for PCA analysis, sample-to-sample Euclidean distance calculations, and heatmaps.

Shrinkage of log fold-changes for volcano plots and Gene Ontology (GO) enrichment analyses was performed on contrasts using apeglm. For each contrast, significantly upregulated and downregulated gene sets (padj < 0.05, |log fold change| > 1) were analyzed separately with clusterProfiler (enrichGO) using the background of all detected genes in the experiment. The ontology category was restricted to Biological Process (BP), p-values were adjusted using Benjamini–Hochberg, and terms with adjusted p < 0.05 (q < 0.2) were considered significant.

### Cell culture

#### ESC maintenance and culture

H9 ESCs were maintained in mTeSR plus media and passaged at least twice post-thaw onto Matrigel coated plates before NMP induction. ESCs were passaged when they reached 70% confluency using ReLeSR and were plated at a 1:4 dilution ratio. The newly passaged hESC cultures reached 70% confluency within 4 days.

#### NMP induction

To induce 2-, 4-, and 10-day NMPs, hESCs were cultured in a N2/B27-based SAND media without vitamin A (IMDM, N-2 1%, B-27 without vitamin A 2%, GlutaMAX 1x, NEAA 1x) under adherent culture conditions. Briefly, NMP induction was initiated by first pulsing the hESCs with 10ng/mL of bFGF for one day followed by the combination of bFGF + Wnt agonist, CHIR 99021. For example, to derive 2-, 4-, or 10-day NMPs, the differentiating hESCs received one day of 10ng/mL bFGF, followed by bFGF + 3μM CHIR 99021 for one day (2-day NMPs), three days (4-day NMPs), or nine days (10-day NMPs) respectively.

#### Embryoid body (EB) differentiation

Upon the completion of NMP induction in adherent cultures, cell sheets containing NMPs were cut with an EZ passage tool and transferred to the ultra-low attachment plates which allowed NMP clumps to self-aggregate into EBs. EBs were cultured with SAND media containing vitamin A (SAND + B27 with vitA) and further supplemented with 1μM RA for two days to induce neural patterning. From the 2nd day until the 14th day, EBs were treated with various patterning factors to induce relevant cell types. For example, to induce dI4-6 populations, EBs were treated with 1μM RA ± GDF11 up to 14 days in suspension cultures. For inducing dI1-3 cell types, EBs were bathed in SAND media supplemented with 1μM RA + 10ng/mL BMP4, and ± GDF11 (50/100ng/mL). On the 14th day EBs were switched into maturation media until day 20, day 43, or day 130 and sampled for quality control checkpoints during maturation. Media changes were performed by collecting EBs in 15mL conical tubes and allowing them to sink to the bottom of the tube in a pellet. Exhausted media was aspirated and replaced with fresh media corresponding to condition. EBs were then transferred back to the plates using glass borosilicate pipettes which prevented EB loss due to sticking to the walls of the plastic pipettes.

#### Immunocytochemistry

NMPs:Immunocytochemistry (IHC) was performed on NMPs to determine the expression of NMP markers such as SOX2 and Brachyury (T). Briefly, at the end of NMP culture durations, the sheet of NMP cells was cut into pieces by EZ Passage tool and transferred onto Matrigel coated 8-well Ibidi slide chambers. In Ibidi slides, NMPs were allowed to grow for another day in their respective condition media and were processed for IHC the following day. NMPs were washed once with 1X DPBS and then fixed on ice with 4% PFA for 8–10 minutes. PFA solution was removed by three 5-minute washes with 1X DPBS, followed by incubation in the antibody blocking solution (1X DPBS, 1% HIHS, 0.1% Triton X-100, and 0.05% sodium azide) for 1 hour at 2–8°C. Primary antibodies were diluted in antibody blocking solution and added to the NMPs for the overnight incubation at 2–8°C. The following dilution was used: SOX2 (mouse; Santa Cruz, sc-365823; 1:500), BRACHYURY (goat; R&D systems, AF2085; 1:500), and CDX2 (rabbit, Cell Signaling Technology, D11D10; 1:500). NMPs were washed twice with 1X DPBS with 0.1% Triton X-100 (PBT) for 5 minutes before secondary antibody incubation. Corresponding species-specific secondary antibodies (Jackson ImmunoResearch Laboratories) were diluted 1:400 in 1X PBT and allowed to incubate in the dark at room temperature for 1 hour followed by counterstaining with nuclei stain DAPI (1:500 dilution in 1X PBT) for 15 minutes and replaced with 1X DPBS for imaging.

Embryoid bodies (EBs): EBs were first collected by the sinking method, and the media was removed by two 5-minute washes with 1X DPBS, followed by fixing in 4% PFA on ice for 10 minutes on rocker. EBs were then washed three times for 5 minutes before allowing them to sink in 30% sucrose in 1X PB solution for at least 1 hour or until all the EBs had sunk to the bottom of the tube, indicating adequate equilibrium with sucrose had been reached. EBs were then embedded in OCT on dry ice blocks, and then sectioned at 14μm thick sections using Leica Cryostat onto positively charged microscope slides. Slides with EB sections were then washed with 1X DPBS, blocked with the antibody blocking solution for 1 hour at room temperature before receiving primary antibodies. The following antibodies were used: LHX2 (mouse; DSHB, PCRP-LHX2-1C11; 1:50), FOXD3 (guinea pig, a gift from Thomas Mueller, Germany, 1:10,000), ISL1 (goat; R&D systems, AF1837; 1:500), PAX2 (rabbit; Invitrogen, 71-6000; 1:500), LMX1B (guinea pig; Thomas Muller Lab), LHX1/5 (mouse; DSHB, 4F2; 1:50) for 12–14 hours (overnight) at 2–8°C in a humidified chamber. Slides were washed twice with PBT before receiving species-specific secondary antibodies (Jackson ImmunoResearch) 1:400 in the dark for 1 hour at 20°C. Slides were then washed once with 1X PBT for 5 minutes and incubated with DAPI for 15 minutes to counterstain the nuclei. Slides were then mounted with Prolong Diamond antifade medium and imaged using a Zeiss LSM800 inverted confocal microscope.

### Quantitative (q) RT-PCR analysis

Samples of interest were lysed with ∼300–500μL RLT buffer and total RNA was purified using Qiagen RNeasy Kit. Total RNA was quantified on a microvolume spectrophotometer, and 500ng/µL RNA was used to generate cDNAs using Superscript IV First Strand Synthesis kit. The gene-specific primers (forward and reverse) were used at 0.5μM final concentration in a 10μL reaction with SYBR Green and qRT-PCR was performed on Roche 480 Light-cycler (Roche). The relative fold expression was determined using the ΔΔCT method to compare the expression of the target gene against the expression of a housekeeping gene glyceraldehyde 3-phosphate dehydrogenase (Gapdh).

## Acknowledgments

We would like to thank Bennett Novitch and members of the Butler and Novitch laboratories for discussions. These studies were supported by a UCLA Broad Stem Cell Research Center (BSCRC) postdoctoral training grant (to S.G._1_) and awards (to S.J.B), the California Institute for Regenerative Medicine (CIRM) Bridges to Research program (EDUC2-12718 to C.R., T.D., T.P., and Y.V.), and a graduate fellowship from the National Institutes of Health (NIH) Training Grant in Genomic Analysis and Interpretation (T32HG002536 to S.G._2_) and grants to S.J.B. from the NIH (R01NS123187) and the Marcus Foundation (4492).

## Author Contributions

S.G._1_, E.H. and S.J.B. conceived the project. E.H. generated the harmonized atlas, SG_1_ and CR led the experiments developing the directed differentiation approaches, with T.D., T.P., A.T. and Y.V. additionally assisting with performing the cellular differentiations and IHC/qPCR analyses.

E.F. analyzed the day 0 bulk and scRNA-Seq NMP and day 27 scRNA-Seq data, S.G._2_ performed the GO analyses on *in vivo*/*in vitro* dI4/dI5s, and E.H. and E.F. both analyzed the day 43 scRNA-Seq data. This paper was written by S.G_1_., E.H., S.G_2_. E.F. and SJB and edited by E.H., S.G._1_ and S.J.B.

---

**Extended Data Figure 1:**
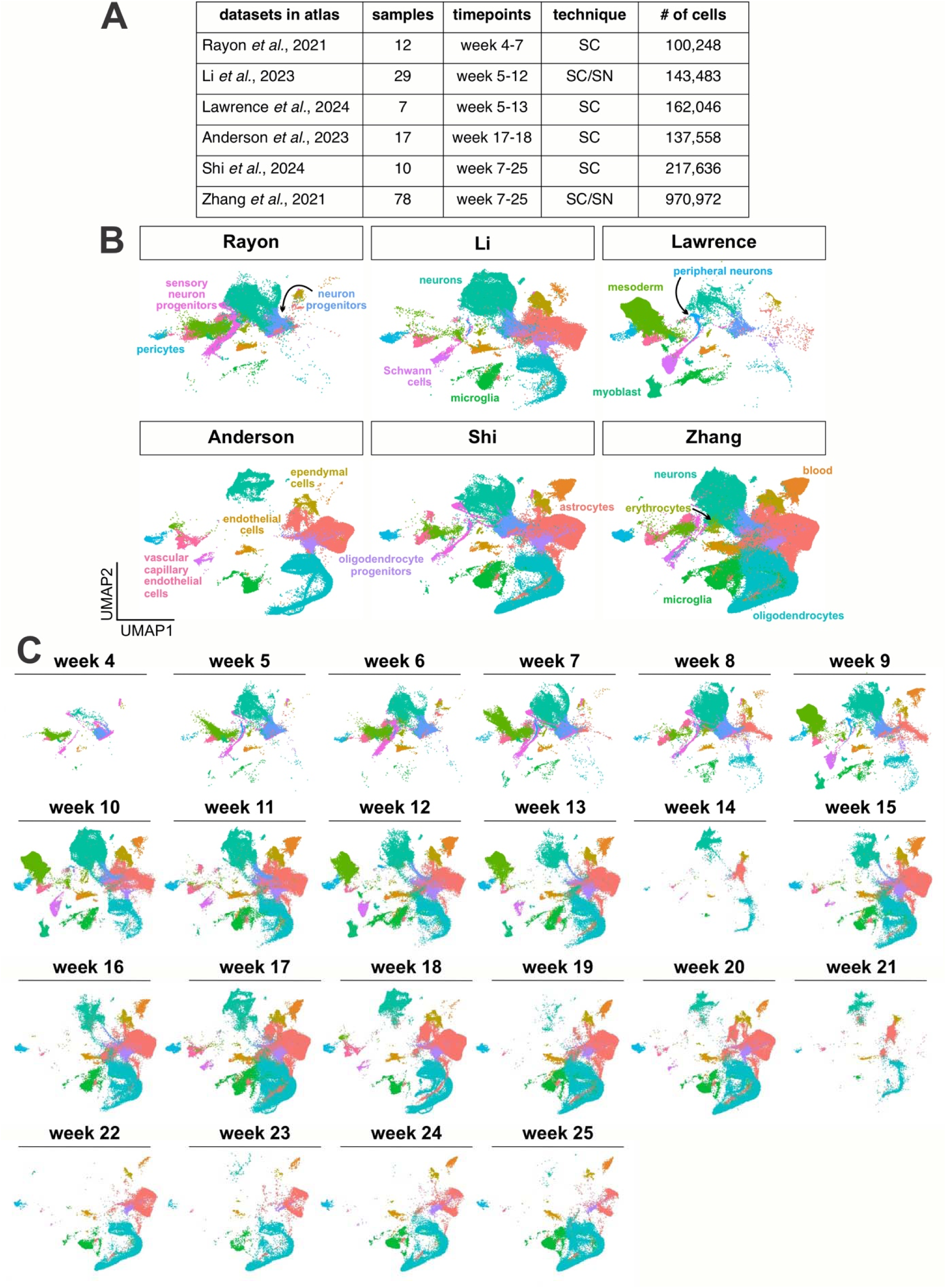
Annotation of different data sets by week in the complete *in vivo* atlas. (A) Table showing summary statistics of the 6 datasets used in generation of the atlas. (B) These datasets each contain different mixtures of cell types, with some containing substantially more neuronal lineage cells than others (C) UMAPs split by week, showing progression of cell type differentiation over time, with early expansion of neurons and mesoderm, followed by astrocytes and oligodendrocytes

**Extended Data Figure 2:**
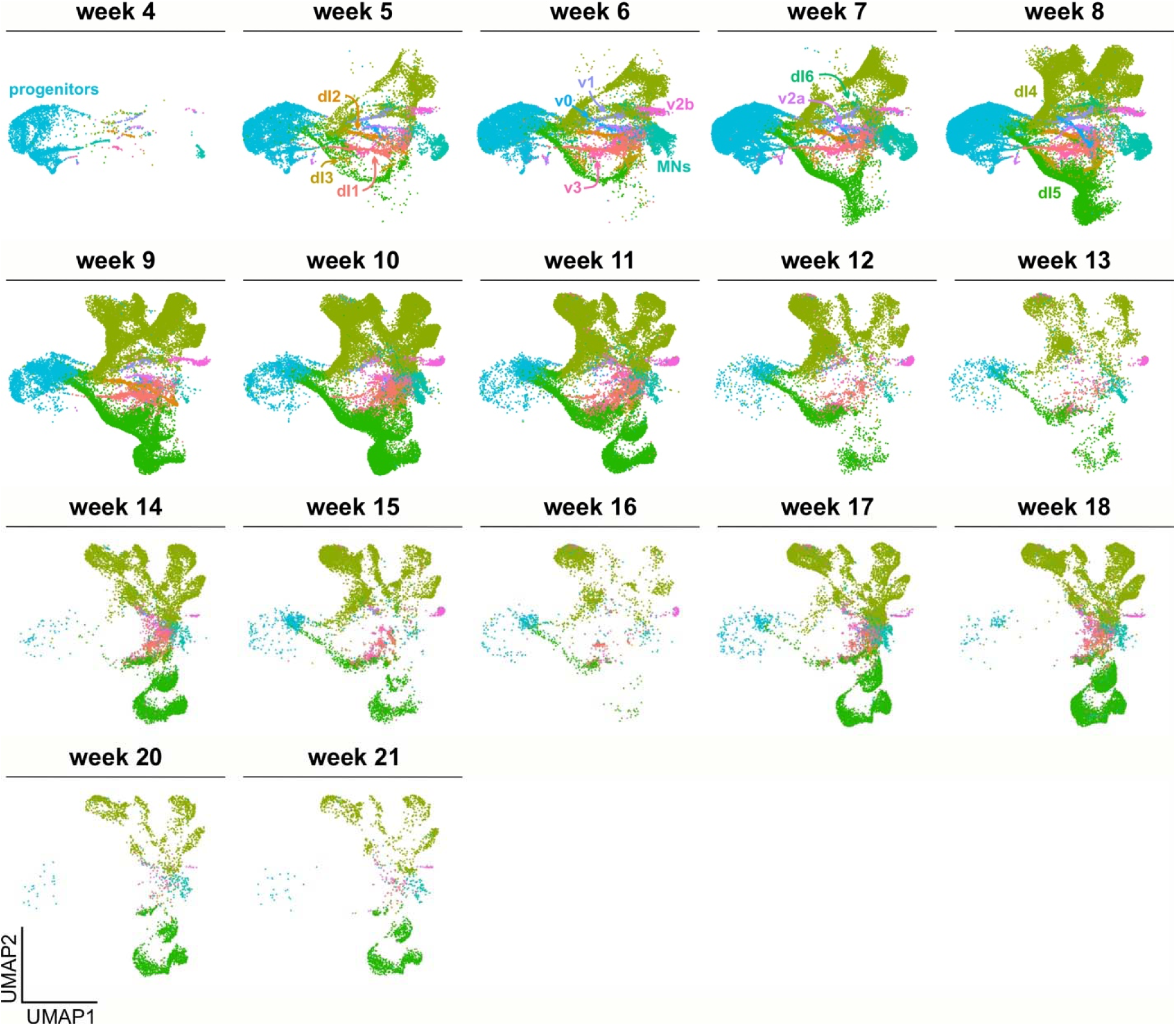
Annotation of cell types in the neuronal *in vivo* atlas by week. Breakout of neuronal atlas cell types by week of sample. Early weeks contain more progenitors and earlier stages of neuronal types. Later weeks contain mostly cells in the extremities of the UMAPs, many of these cell types are cells that contain gene expression overlapping with dILs.

**Extended Data Figure 3:**
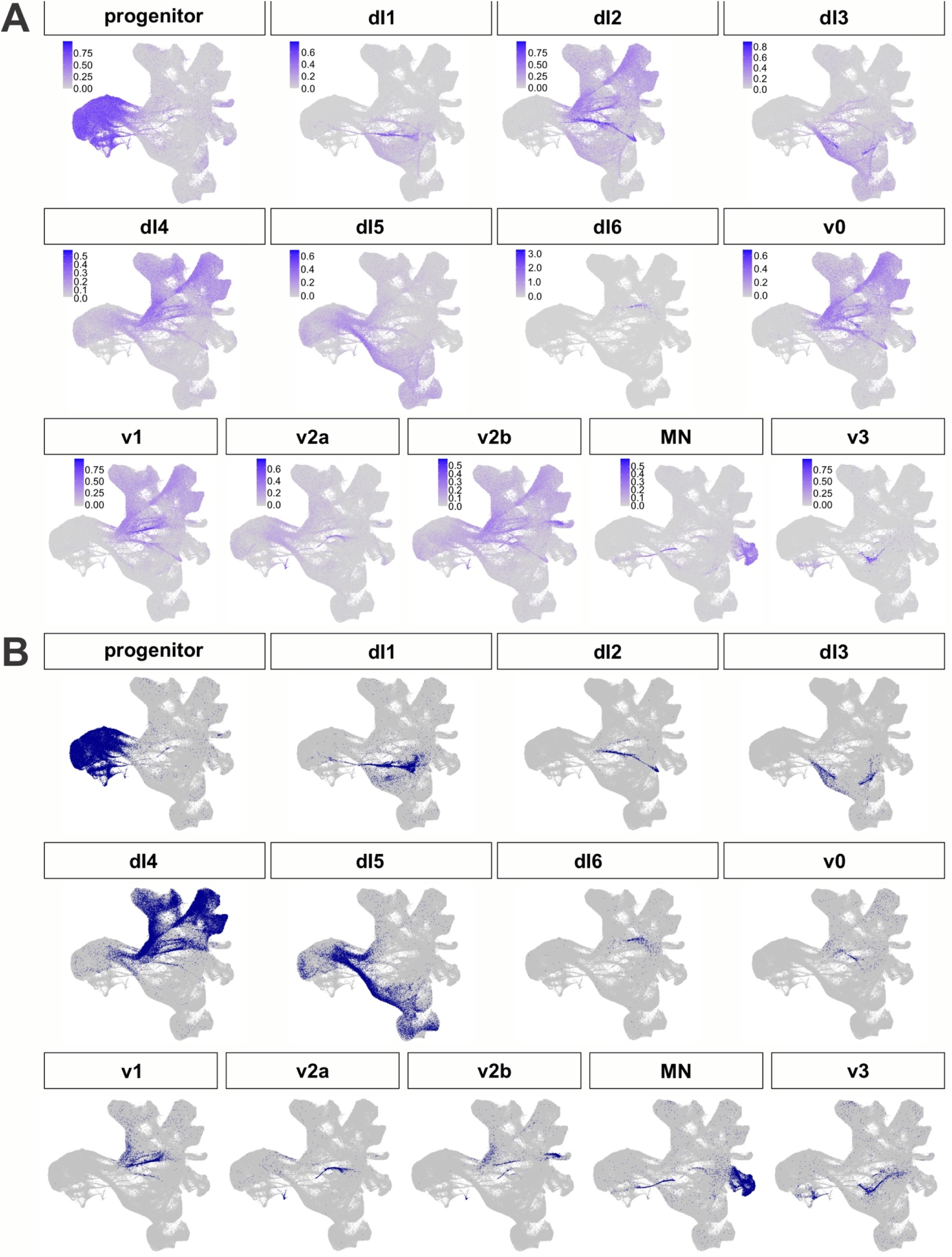
Assignment of spinal trajectories within the *in vivo* atlas. (A) AUCell was used with a custom marker gene list to plot module scores for each neuronal lineage in the atlas. For dI6, expression of DMRT3 is shown since AUCell did not provide sufficiently specific scores. (B) Cutoffs were manually assigned for each cell type to determine which cells could be classified as each cell type. Cells with multiple identities were then rectified with kNN analysis, and then identities were extrapolated to cells with no identity using kNN analysis as well.

**Extended Data Figure 4:**
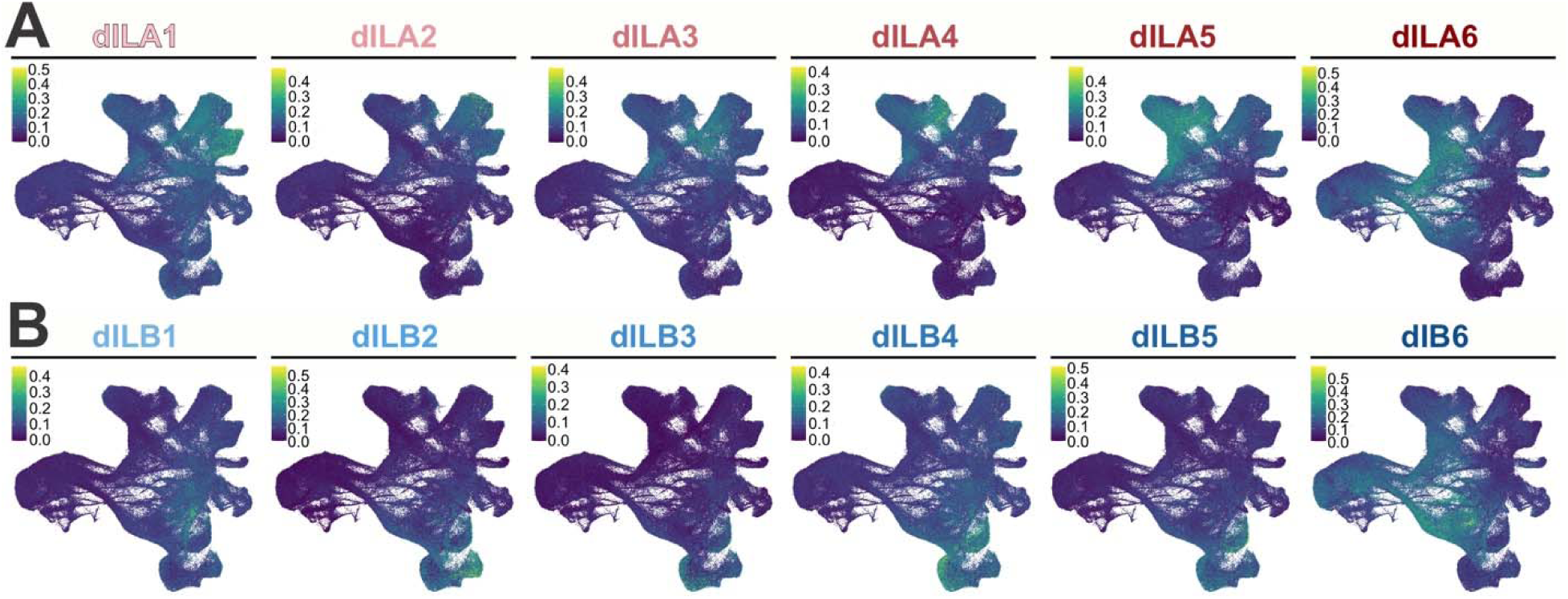
Assignment of mouse dI4/dI5 identities on human neural atlas. (A-B) Gene lists from Roome et al^50^ of each dIL subtype were used in AUCell to calculate module scores for each of the potential dIL subtypes. This was then used to map their general location onto our atlas.

**Extended Data Figure 5:**
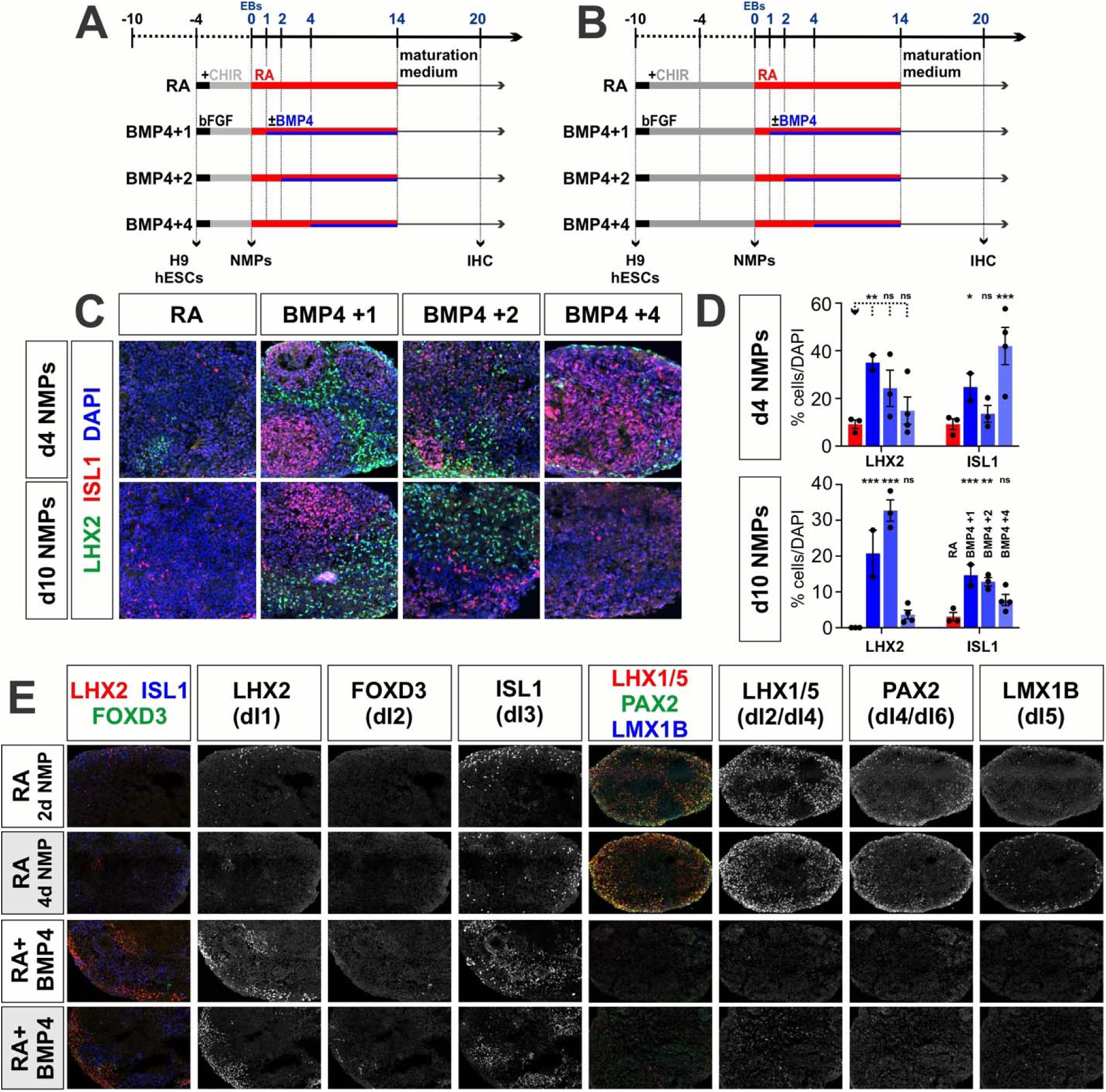
Assessing the timing of BMP4 addition in d4 and d10 NMP protocols. (A, B) EBs were generated at day 0 after either d4 (A) or d10 (B) NMP formation and then treated with RA for 10 days (control). In the experimental samples, BMPs were added at day 1 (BMP4+1), day 2 (BMP4+2) or day 4 (BMP4+4) after EB formation, until day 10. Samples were further differentiated for 6 days and then processed for immunohistochemistry (IHC). (C) EBs were labeled with antibodies against LHX2 (dI1, green), FOXD3 (dI2, white), ISL1 (dI3, red) and DAPI (all nuclei, blue). (D, E) Quantification of EBs formed from either d4 (D) or d10 (E) NMPs, suggests that dI1s form robustly from the BMP4+1 condition for both d4 and d10 NMPs. In contrast, dI3s form most robustly from BMP4+4 condition for d4 NMPs and the BMP4+1 condition for d10 NMPs. Probability of similarity between control and experimental groups: * p<0.05, ** p<0.005, *** p<0.0005; two-way ANOVA.

**Extended Data Figure 6:**
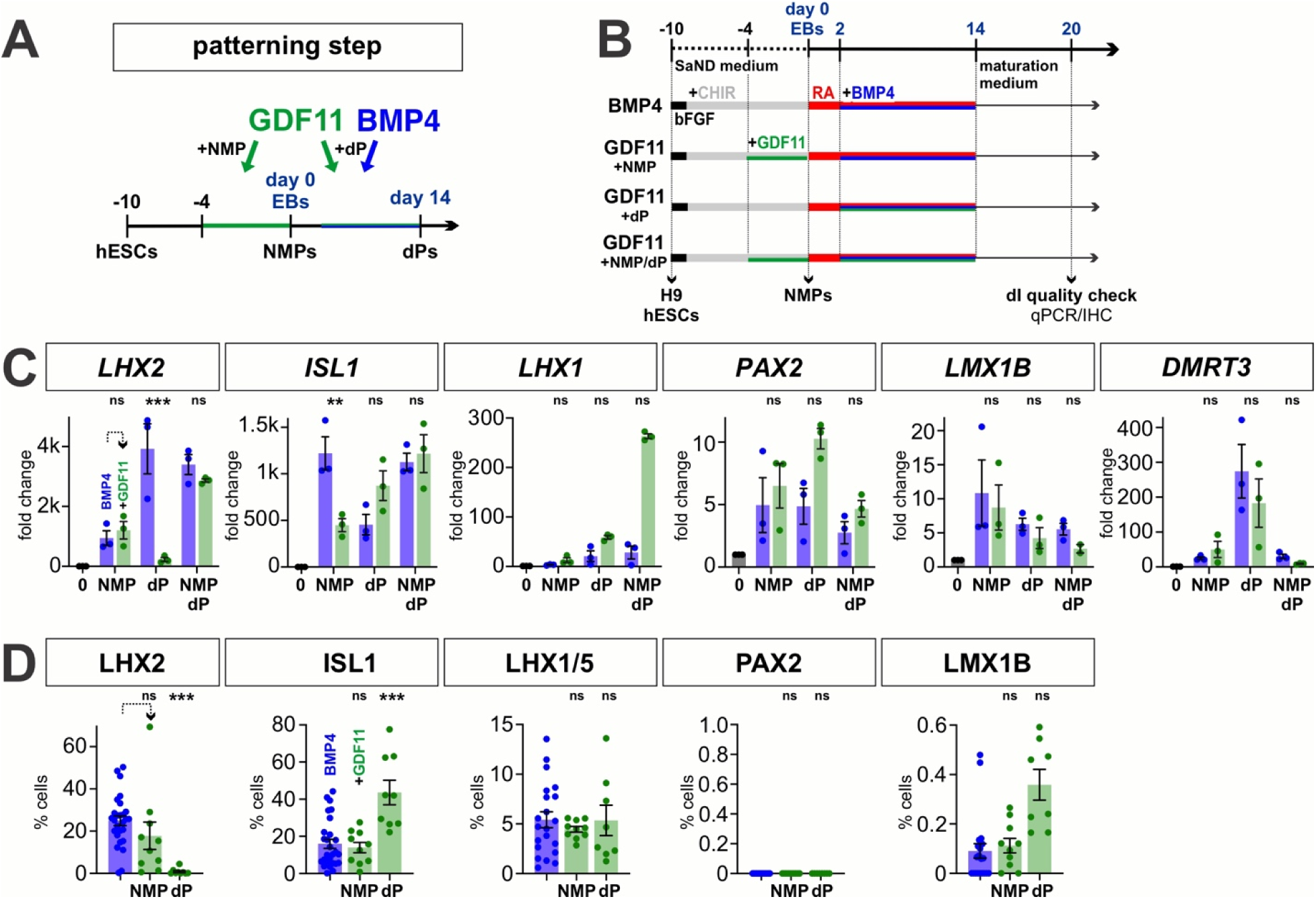
Effect of GDF11 on dI identity in the RA+BMP4 protocol. (A, B) Overview of the experimental timeline/workflow for the RA±BMP4±GDF11 NMP protocols. The effect of GDF11 treatment was assessed on NMP patterning, dP patterning and both NMP and dP patterning (A). (C, D) qPCR (D) and IHC (D) analyses of day 20 EBs, suggest that the addition of GDF11 together with BMP4 at the dP stage significantly decreases LHX2^+^ dI1 identity, but increases ISL1^+^ dI3 identities. There are no other significant effects. Probability of similarity between control and experimental groups: **p<0.005 *** p<0.0005.

**Extended Data Figure 7:**
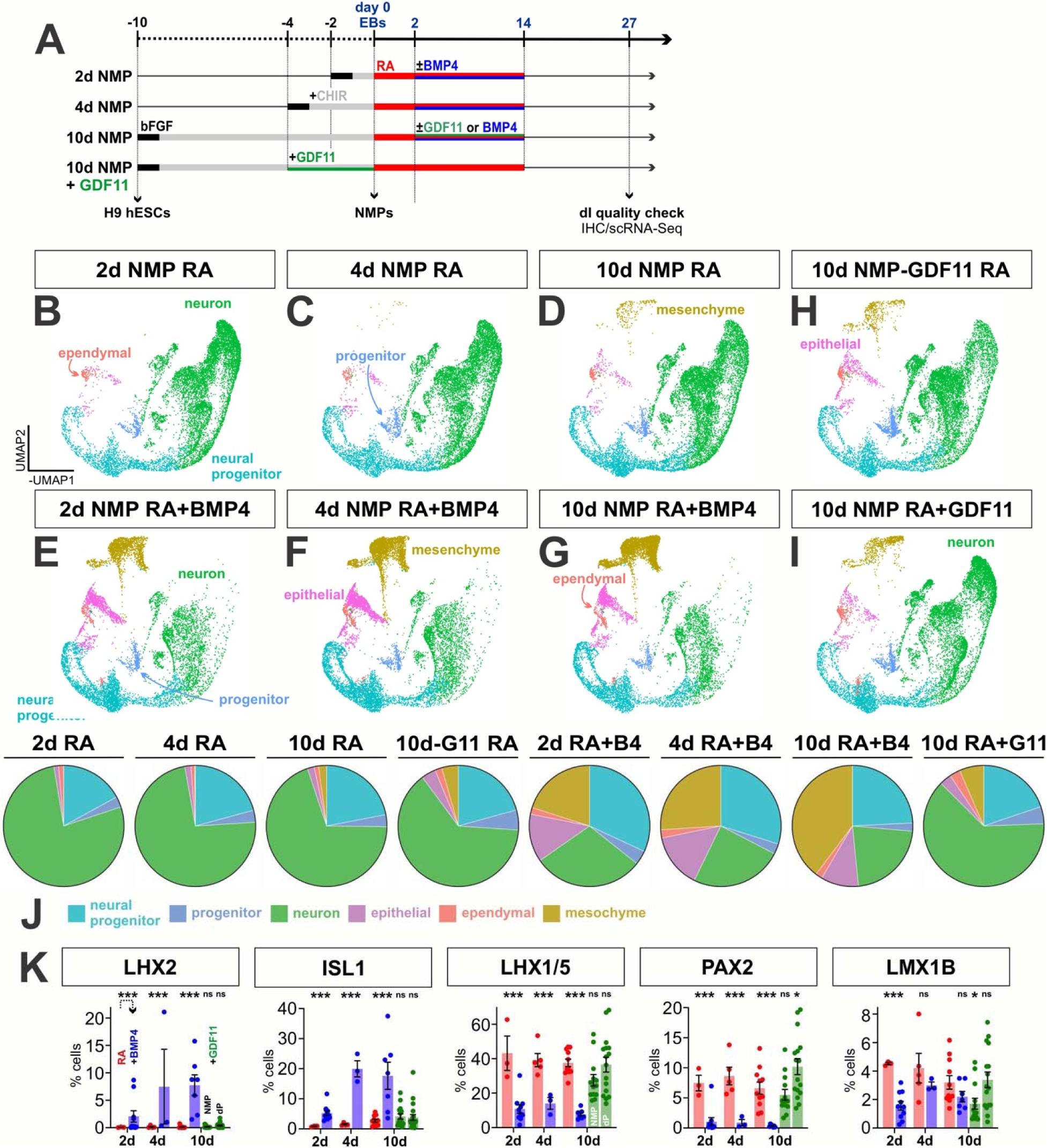
scRNA-Seq analysis of day 27 EBs derived from all major RA, BMP4 and GDF11 conditions. (A) Overview of the experimental timeline/workflow for this set of RA±BMP4±GDF11 NMP protocols. ScRNA-Seq and IHC data was analyzed at day 27. (B-J) UMAPs of days 27 in vitro-derived cells generated using the RA, RA+BMP4, and RA+GDF11 (NMP and dP) protocols, showing the automated celltype identity assignment on a cell-by-cell basis using the curated marker genes shown in Fig. 1d. The pie charts (J) summarize the proportions of cells generated by each protocol. The 2d, 4d,10d NMP cultures performed equivalently in the RA protocol, proficiently generating neuronal cell types (>90%). The addition of GDF11 at either the NMP or dP stage result in ∼80% neuronal cells, while the addition of BMP4 results in both neural cells (∼50% neuronal, ∼7% glia), and other ectodermal (skin, ∼12%) and mesodermal (∼28%) derivatives. The BMP-dependent non-neuronal derivatives increase, as NMPs spend more time in culture. (K) IHC analyses of day 27 EBs, reproduce the expected result that the addition of BMP4 increases the numbers of the LHX2^+^ and ISL1^+^ dorsal most dIs, while the RA alone condition directs the intermediate PAX2+ and LMX1B+ dIs. In this experiment, GDF11 added as a dP patterning factor, significantly increased the PAX2^+^ identities, with only a trend towards increasing LMX1B^+^ identities. Probability of similarity between control and experimental groups: **p<0.005 *** p<0.0005.

**Extended Data Figure 8:**
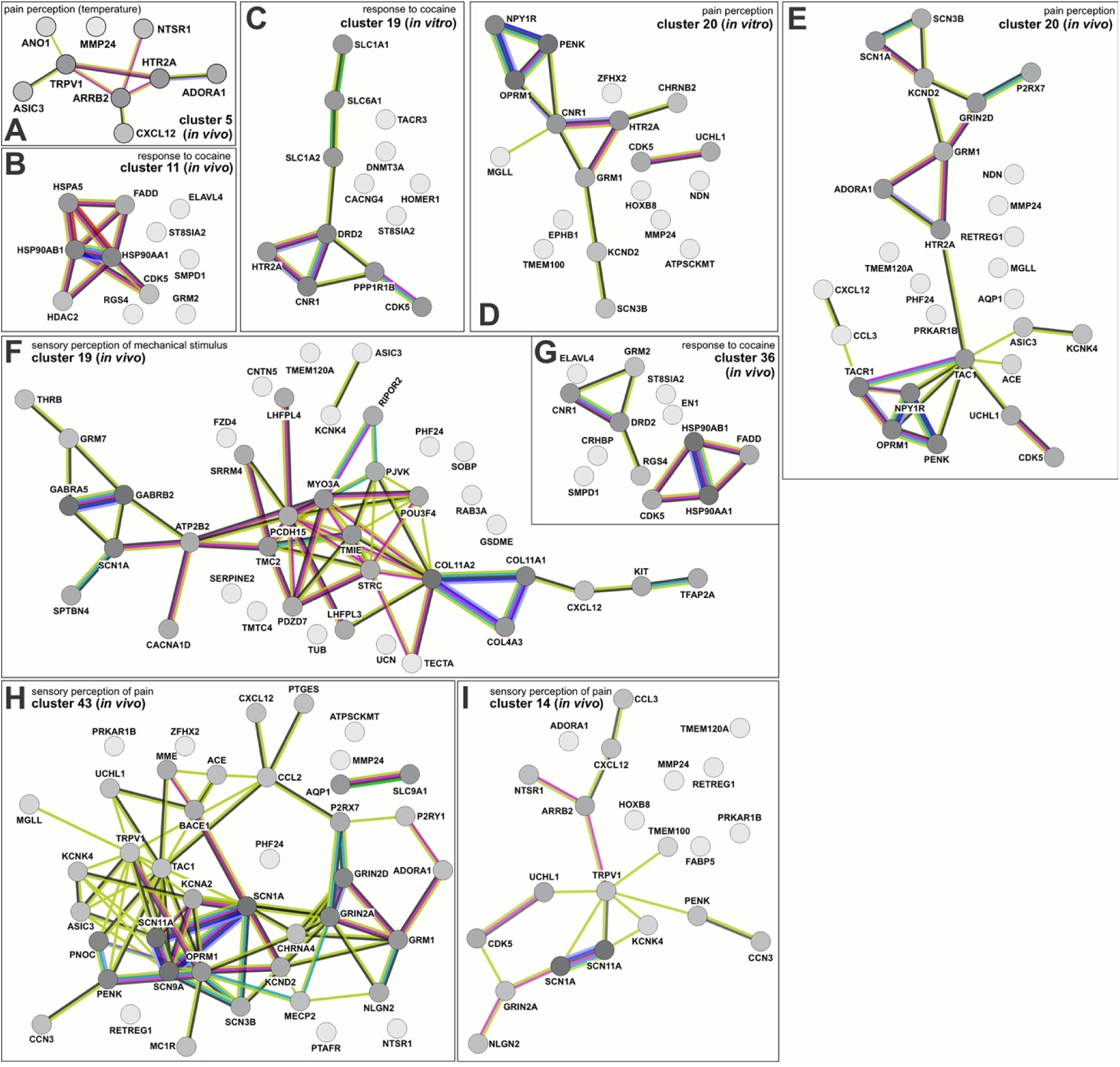
Pain perception, cocaine response, and mechanosensory networks are found in *in vivo* and *in vitro* dI4 clusters. The STRING (search tool for the retrieval of interacting genes/proteins)^59^ was used to identify sensory modality networks from genes upregulated in both *in vivo* and *in vitro* dI cell populations. Nodes represent proteins, the lines (or edges) between the nodes represent protein-protein associations. Numerous edges between nodes implies greater evidence (experimentally determined or predicted) for protein-protein associations. STRING further documents evidence of protein associations through the edge color as follows: Known interactions: experimentally determined (purple) or from curated databases (teal). Predicted interactions: gene neighborhood (green), gene fusions (red), gene co-occurrence (blue). Others: text mining (yellow), co-expression (black), protein homology (periwinkle). (A, D, E, H, I) Networks associated with pain perception (A) *In-vivo* cluster 5 cells make up the sole network linking temperature with pain, centering around NTSR1, TRPV1, ARRB2 AND HTR2A. Pain circuits regulated by PENK, OPRM1, NPY1R, HTR2A and GRM1 are found in both *in vitro* cluster 20 (D) and *in vivo* cluster 20 (E). The *in vivo* cluster 43 (H) circuit has similarity with *in vivo* cluster 14 (I), sharing the SCN1A, SCN11A, and TRPV1 nodes. (B, C, G) Protein networks involved in response to cocaine. Both *in vitro* cluster 19 (C) and *in vivo* cluster 36 (G) circuits center around DRD2 and CNR1, further evidence supporting our *in vitro* generated dI4 cells may be functionally similar to their *in vivo* counterparts. (F) Sensory perception of mechanical stimulus protein network stemming from upregulated genes found in *in vivo* cluster 19

**Extended Data Figure 9:**
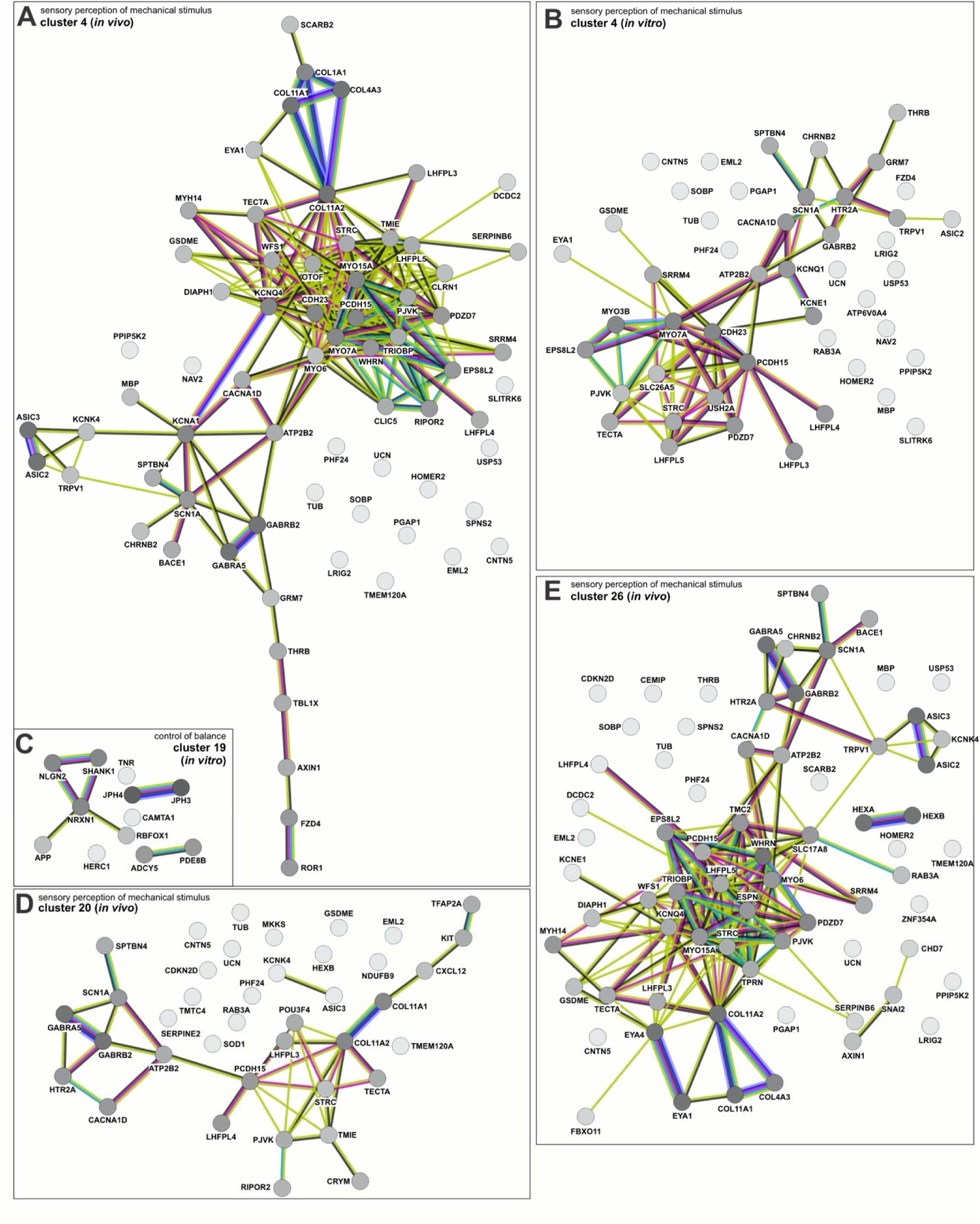
Mechanosensation and balance networks are found among *in vivo* and *in vitro* dI4 clusters. (A, B, D, E) Numerous *in vitro* and *in vivo* clusters contain protein interaction networks involved in mechanosensation. The *in vitro* cluster 4 (B) circuit includes MYO7A, PCDH15, MYO3B and EPS8L2, key nodes also found across *in vivo* clusters 4 (A), 20 (D) and 26 (E). (C) A protein network associated with balance is found in *in vitro* cluster 19. Notably, this includes SHANK1, NLGN2, and NRXN1, genes with allele variants associated with autism spectrum disorder.

**Extended Data Figure 10:**
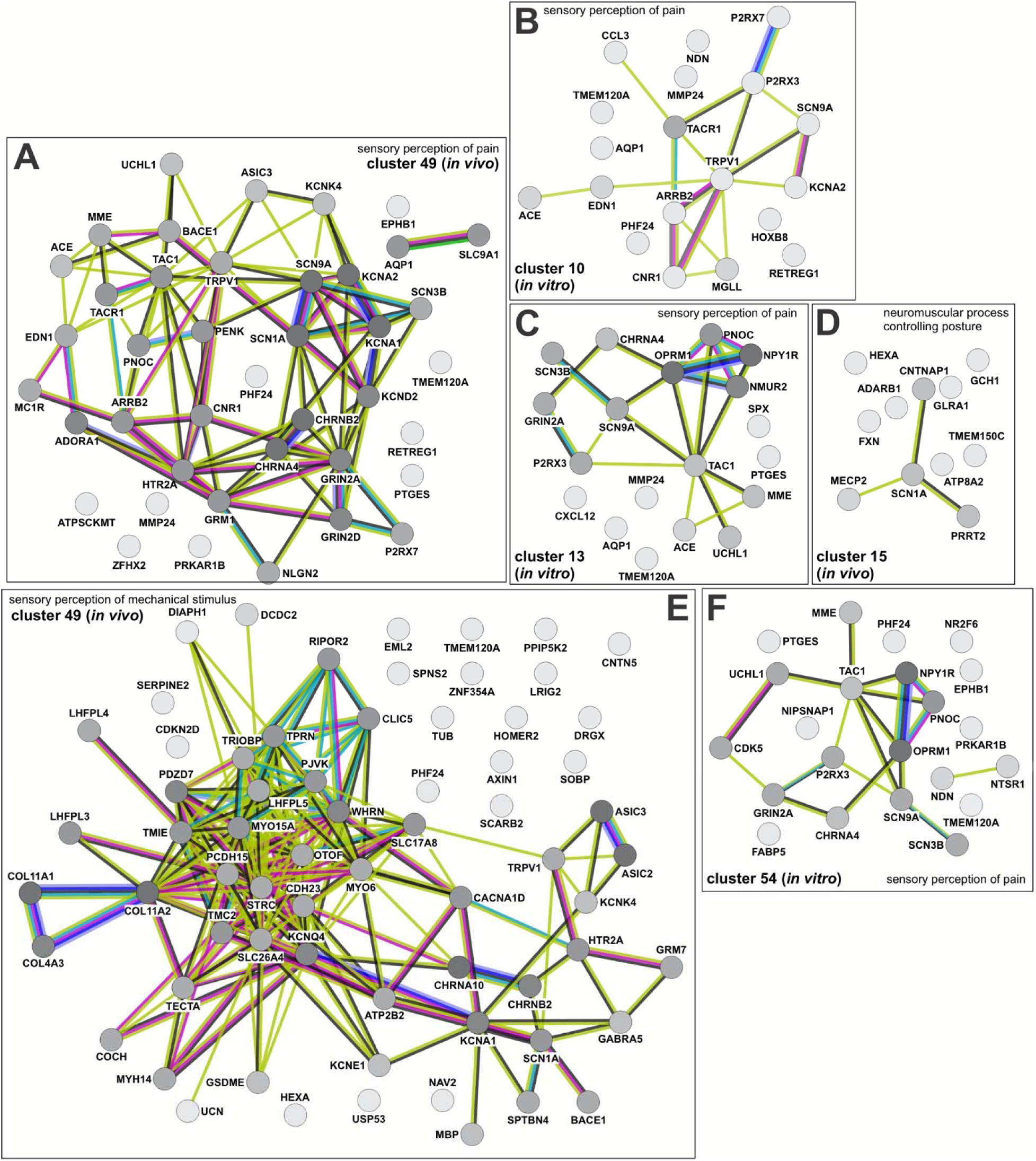
Pain perception, posture regulation, and mechanosensory networks are found among *in vivo* and *in vitro* dI5 clusters. (A-C, F) Networks identified associated with pain perception. The cluster 49 *in vivo* circuit includes strong associations between SCN9A, SCN1A, KCNA1, and SCN3B. Part of this module is recapitulated in *in vitro* cluster 10 (B), with the presence of SCN9A and KCNA2 as well as *in vitro* cluster 54 (F) with the presence of SCN9A and SCN3B. *In vitro* cluster 13 (C) interestingly includes NMUR2, a protein crucial in sensing mechanical itch, which is not present in other identified networks. (D) Unique to the *in vivo* dataset is a neuromuscular posture regulating network comprised of CNTNAP1, SCN1A, PRRT2 and MECP2. (E) A large *in vivo* network associated with mechanosensation is found in cluster 49 and includes similar nodes such as PCDH15 described in the clusters in Extended data figure 8 relating to mechanosensation.

