## Supplemental Figures for "*In vivo* human embryonic spinal cord atlas validates stem cell–derived human dorsal interneurons and reveals ASD spinal signatures"

**Extended Data Figure 1:** Annotation of different data sets by week in the complete *in vivo* atlas

A

| datasets in atlas | samples | timepoints | technique | # of cells |
|---|---|---|---|---|
| Rayon *et al.*, 2021 | 12 | week 4-7 | SC | 100,248 |
| Li *et al.*, 2023 | 29 | week 5-12 | SC/SN | 143,483 |
| Lawrence *et al.*, 2024 | 7 | week 5-13 | SC | 162,046 |
| Anderson *et al.*, 2023 | 17 | week 17-18 | SC | 137,558 |
| Shi *et al.*, 2024 | 10 | week 7-25 | SC | 217,636 |
| Zhang *et al.*, 2021 | 78 | week 7-25 | SC/SN | 970,972 |

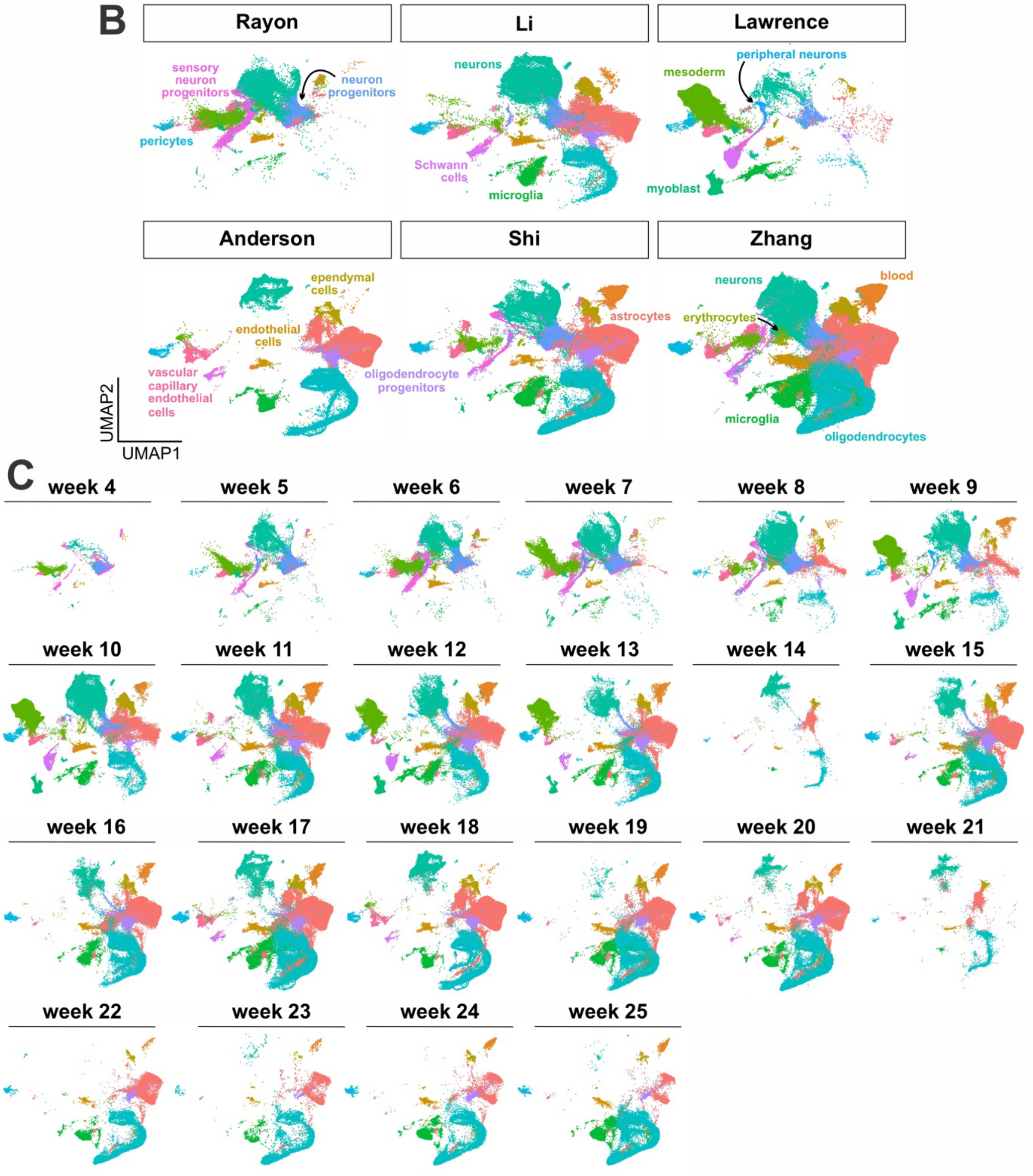

(A) Table showing summary statistics of the 6 datasets used in generation of the atlas.

(B) These datasets each contain different mixtures of cell types, with some containing substantially more neuronal lineage cells than others

(C) UMAPs split by week, showing progression of cell type differentiation over time, with early expansion of neurons and mesoderm, followed by astrocytes and oligodendrocytes

**Extended Data Figure 2:** Annotation of cell types in the neuronal *in vivo* atlas by week

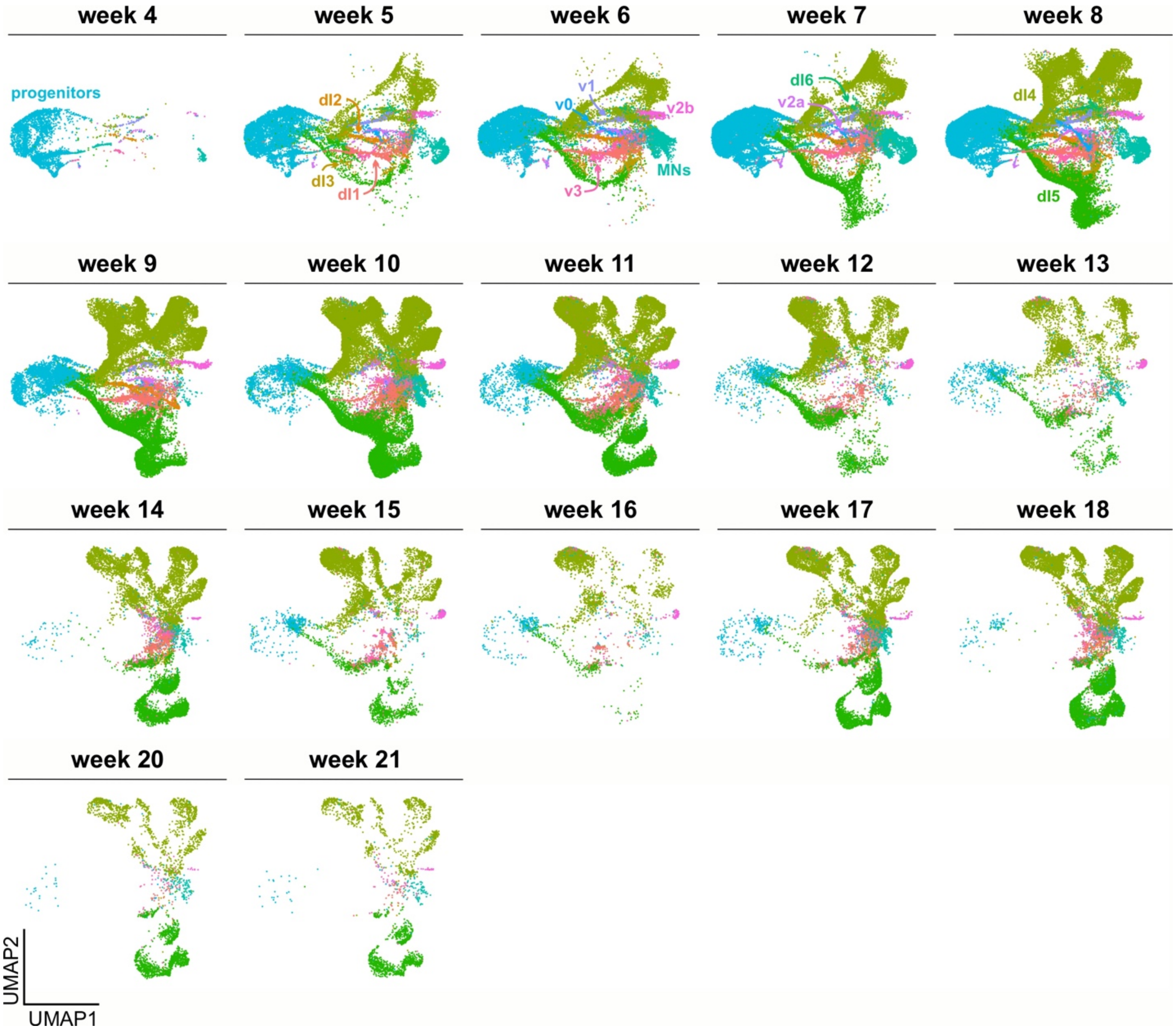


Breakout of neuronal atlas cell types by week of sample. Early weeks contain more progenitors and earlier stages of neuronal types. Later weeks contain mostly cells in the extremities of the UMAPs, many of these cell types are cells that contain gene expression overlapping with dILs.

**Extended Data Figure 3:** Assignment of spinal trajectories within the *in vivo* atlas

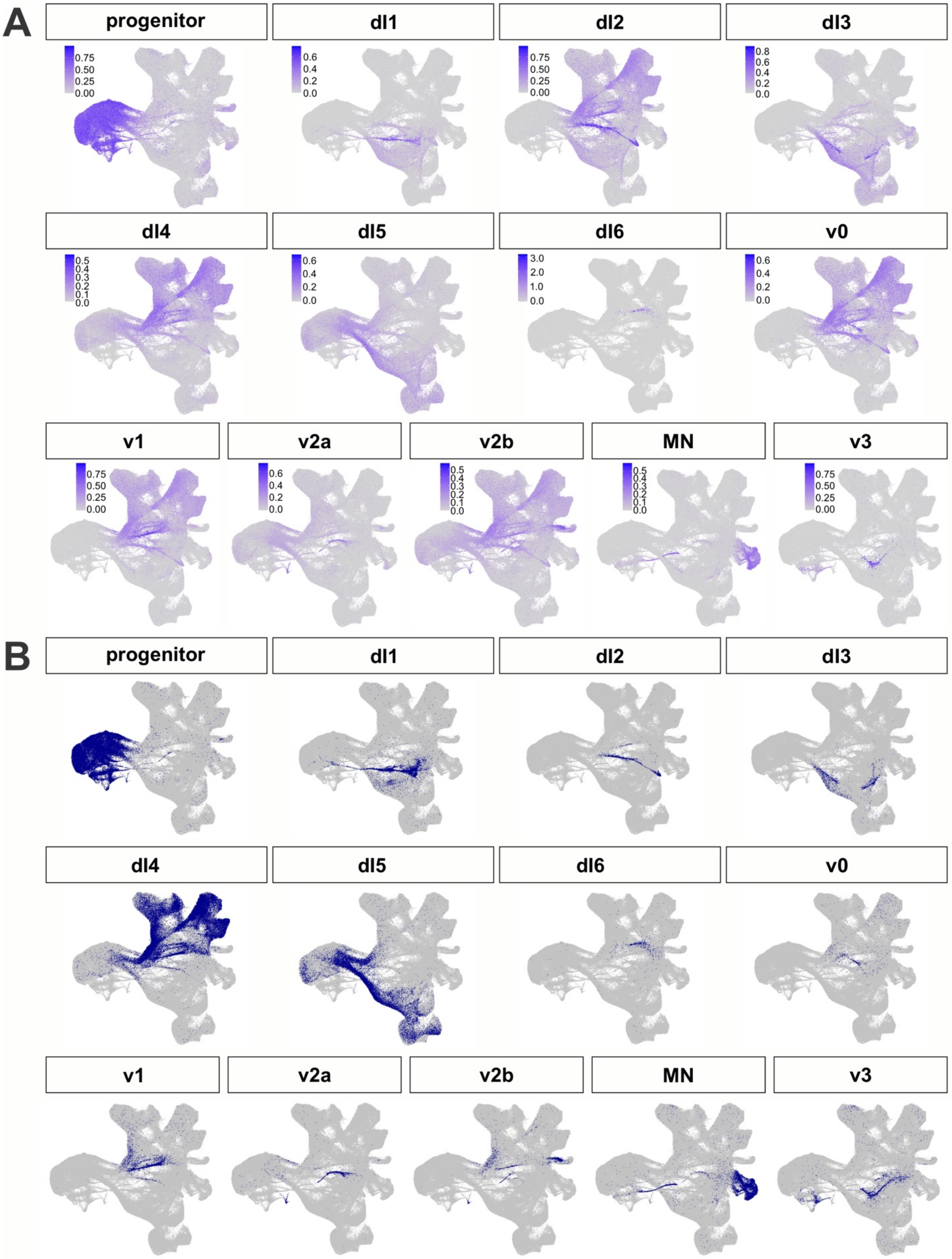

(A) AUCell was used with a custom marker gene list to plot module scores for each neuronal lineage in the atlas. For dI6, expression of DMRT3 is shown since AUCell did not provide sufficiently specific scores.

(B) Cutoffs were manually assigned for each cell type to determine which cells could be classified as each cell type. Cells with multiple identities were then rectified with kNN analysis, and then identities were extrapolated to cells with no identity using kNN analysis as well.

**Extended Data Figure 4:** Assignment of mouse dI4/dI5 identities on human neural atlas

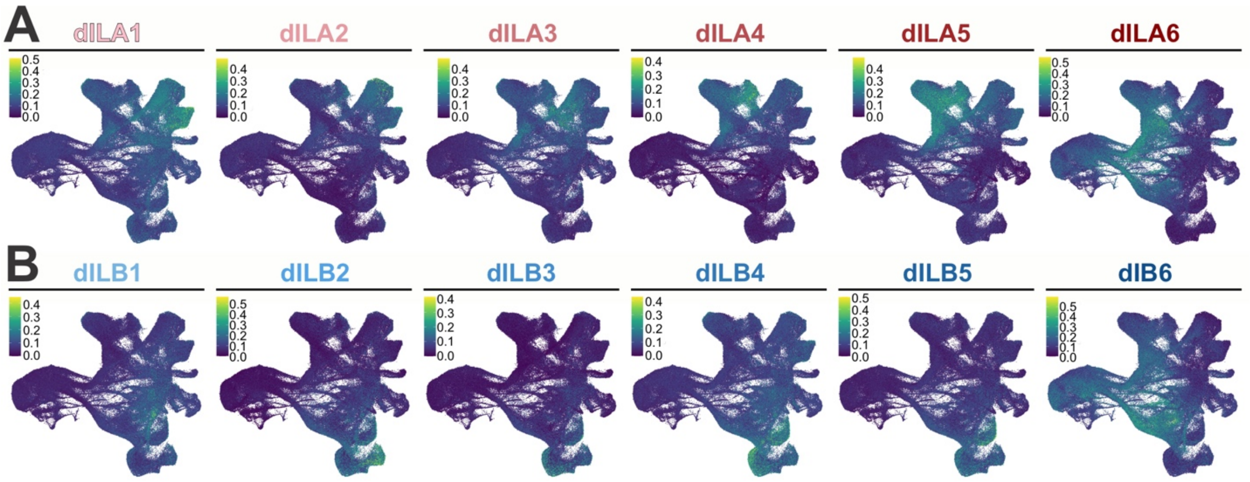


(A-B) Gene lists from Roome et al[50] of each dIL subtype were used in AUCell to calculate module scores for each of the potential dIL subtypes. This was then used to map their general location onto our atlas.

**Extended Data Figure 5:** Assessing the timing of BMP4 addition in d4 and d10 NMP protocols.

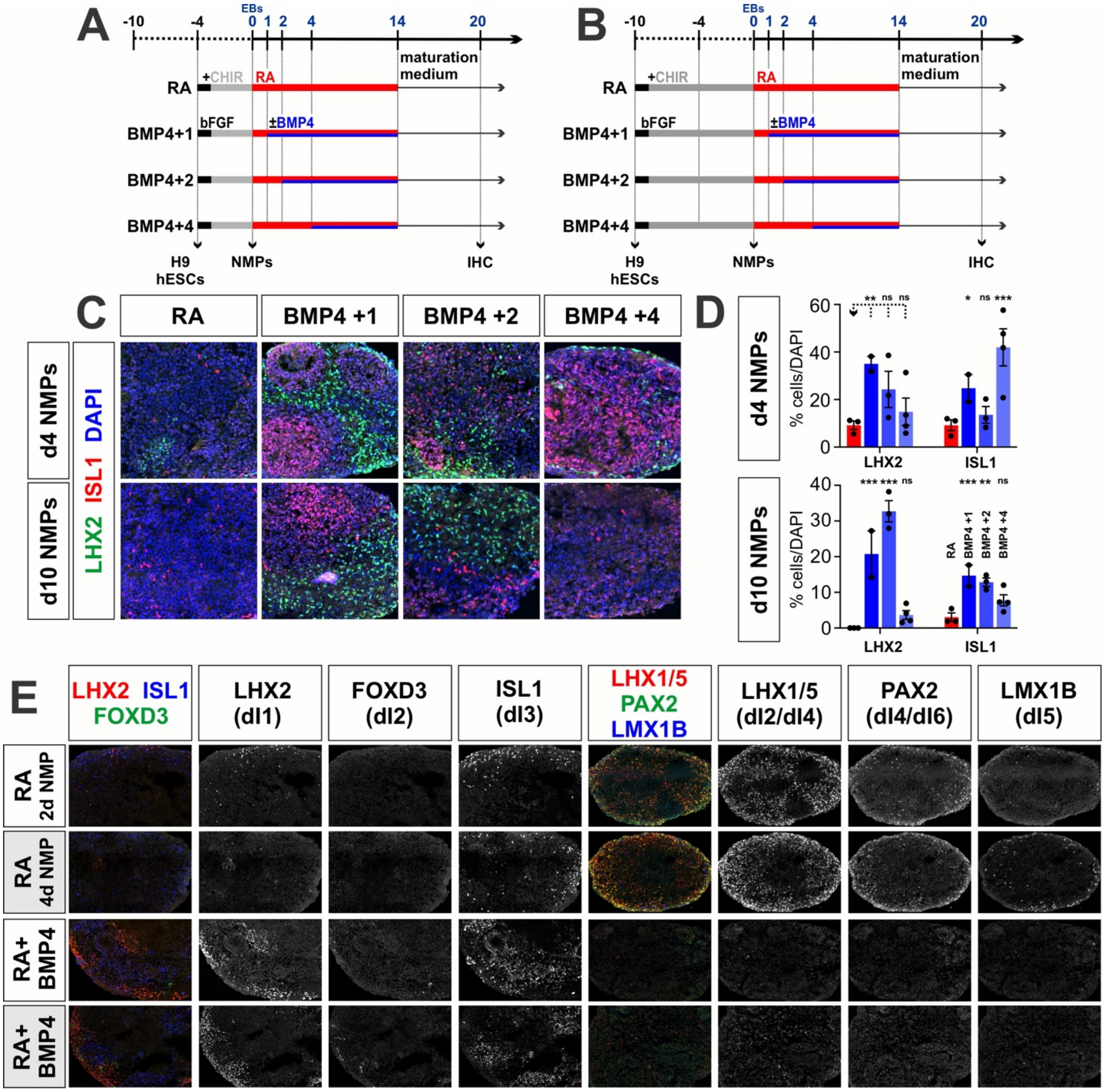

(A, B) EBs were generated at day 0 after either d4 (A) or d10 (B) NMP formation and then treated with RA for 10 days (control). In the experimental samples, BMPs were added at day 1 (BMP4+1), day 2 (BMP4+2) or day 4 (BMP4+4) after EB formation, until day 10. Samples were further differentiated for 6 days and then processed for immunohistochemistry (IHC).

(C) EBs were labeled with antibodies against LHX2 (dI1, green), FOXD3 (dI2, white), ISL1 (dI3, red) and DAPI (all nuclei, blue).

(D, E) Quantification of EBs formed from either d4 (D) or d10 (E) NMPs, suggests that dI1s form robustly from the BMP4+1 condition for both d4 and d10 NMPs. In contrast, dI3s form most robustly from BMP4+4 condition for d4 NMPs and the BMP4+1 condition for d10 NMPs.

Probability of similarity between control and experimental groups: * $p<0.05$, ** $p<0.005$, *** $p<0.0005$; two-way ANOVA.

**Extended Data Figure 6:** Effect of GDF11 on dI identity in the RA+BMP4 protocol

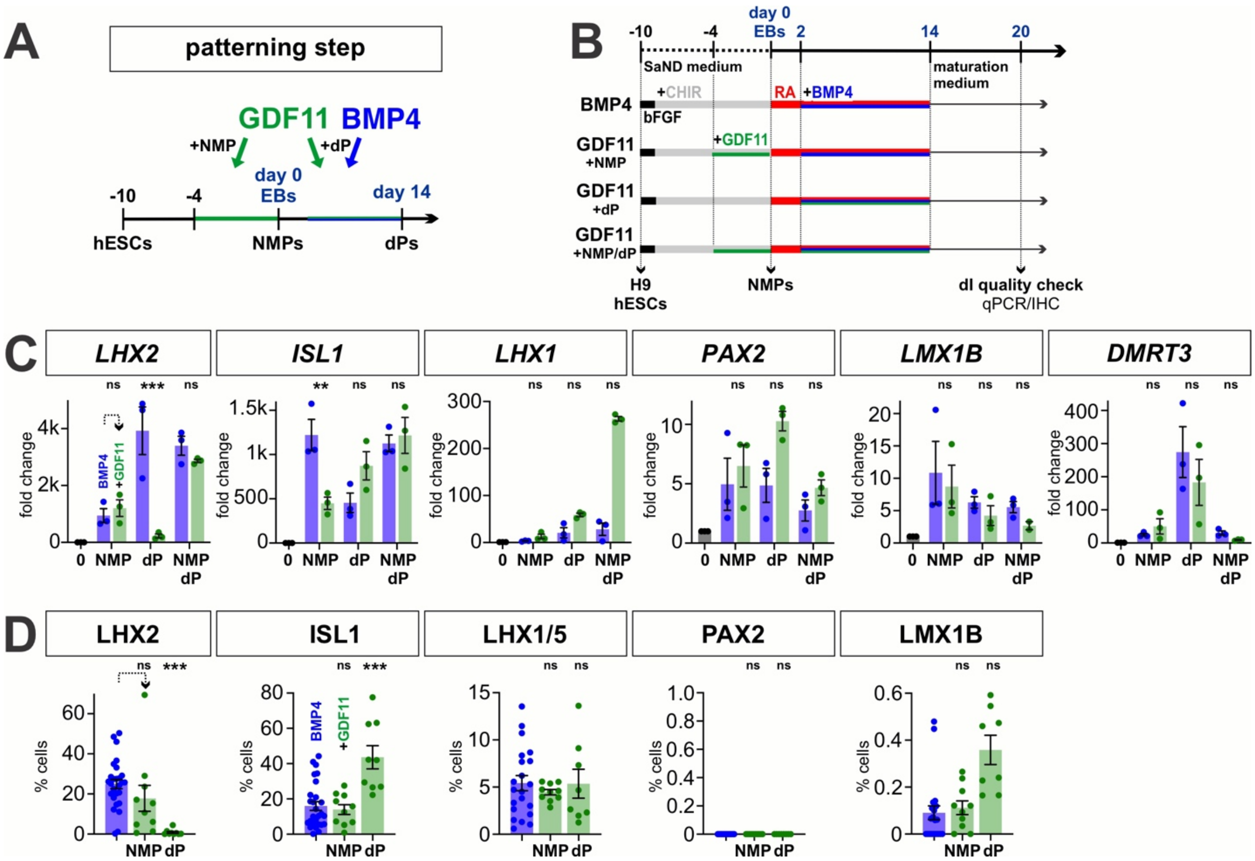


(A, B) Overview of the experimental timeline/workflow for the RA±BMP4±GDF11 NMP protocols. The effect of GDF11 treatment was assessed on NMP patterning, dP patterning and both NMP and dP patterning (A).

(C, D) qPCR (D) and IHC (D) analyses of day 20 EBs, suggest that the addition of GDF11 together with BMP4 at the dP stage significantly decreases LHX2$^{+}$ dI1 identity, but increases ISL1$^{+}$ dI3 identities. There are no other significant effects.

Probability of similarity between control and experimental groups: **p<0.005 *** p<0.0005.

**Extended Data Figure 7:** scRNA-Seq analysis of day 27 EBs derived from all major RA, BMP4 and GDF11 conditions

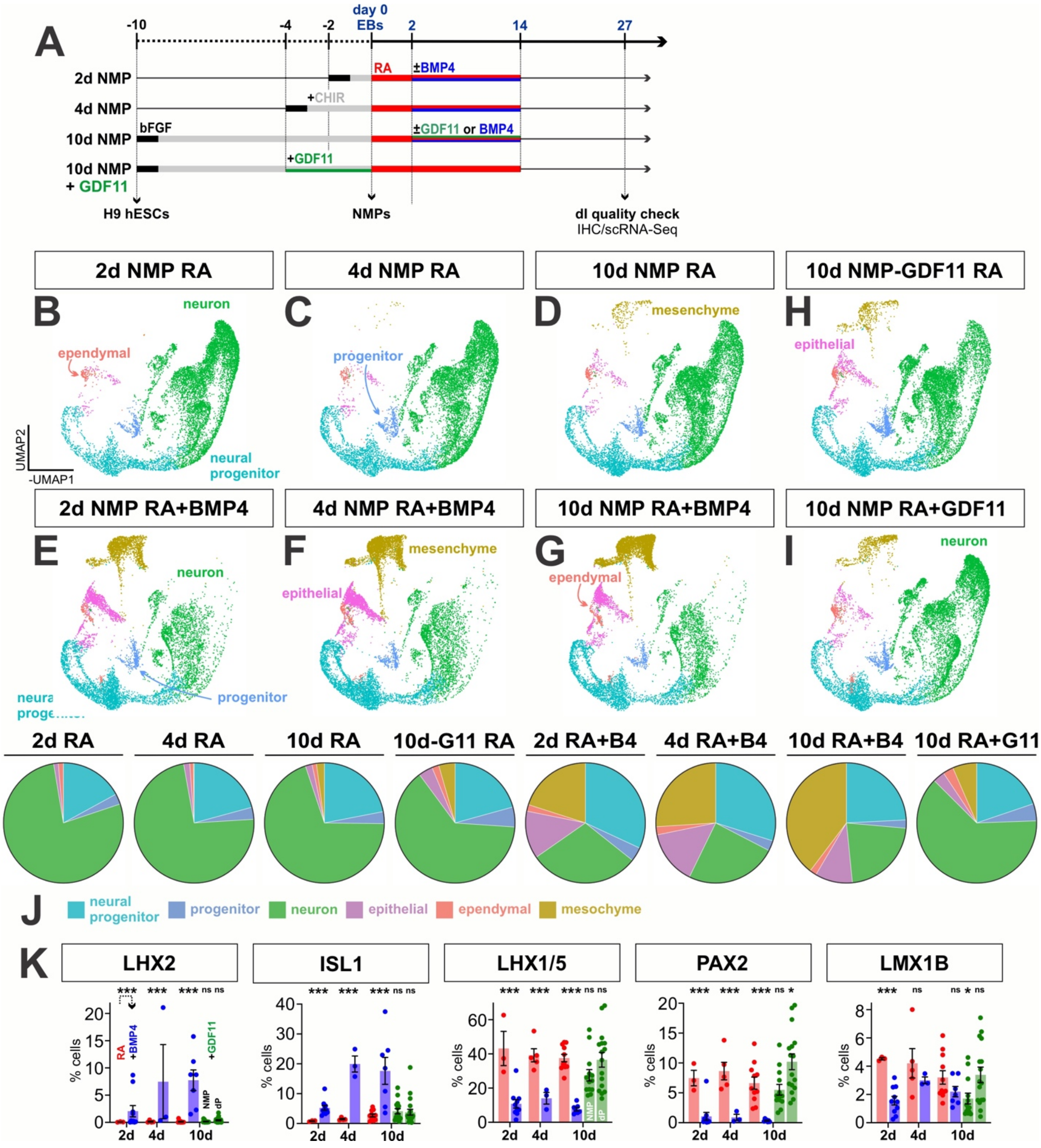


(A) Overview of the experimental timeline/workflow for this set of RA±BMP4±GDF11 NMP protocols. ScRNA-Seq and IHC data was analyzed at day 27.

(B-J) UMAPs of days 27 in vitro-derived cells generated using the RA, RA+BMP4, and RA+GDF11 (NMP and dP) protocols, showing the automated celltype identity assignment on a cell-by-cell basis using the curated marker genes shown in Fig. 1d.  The pie charts (J) summarize the proportions of cells generated by each protocol.  The 2d, 4d ,10d NMP cultures performed equivalently in the RA protocol, proficiently generating neuronal cell types (>90%).  The addition of GDF11 at either the NMP or dP stage result in ~80% neuronal cells, while the addition of BMP4 results in both neural cells (~50% neuronal, ~7% glia), and other ectodermal (skin, ~12%) and mesodermal (~28%) derivatives.   The BMP-dependent non-neuronal derivatives increase, as NMPs spend more time in culture.

(K) IHC analyses of day 27 EBs, reproduce the expected result that the addition of BMP4 increases the numbers of the $LHX2^{+}$ and $ISL1^{+}$ dorsal most dIs, while the RA alone condition directs the intermediate PAX2+ and LMX1B+ dIs.  In this experiment, GDF11 added as a dP patterning factor, significantly increased the $PAX2^{+}$ identities, with only a trend towards increasing $LMX1B^{+}$ identities.

Probability of similarity between control and experimental groups:  **$p<0.005$ *** $p<0.0005$.

**Extended Data Figure 8:** Pain perception, cocaine response, and mechanosensory networks are found in *in vivo* and *in vitro* dI4 clusters

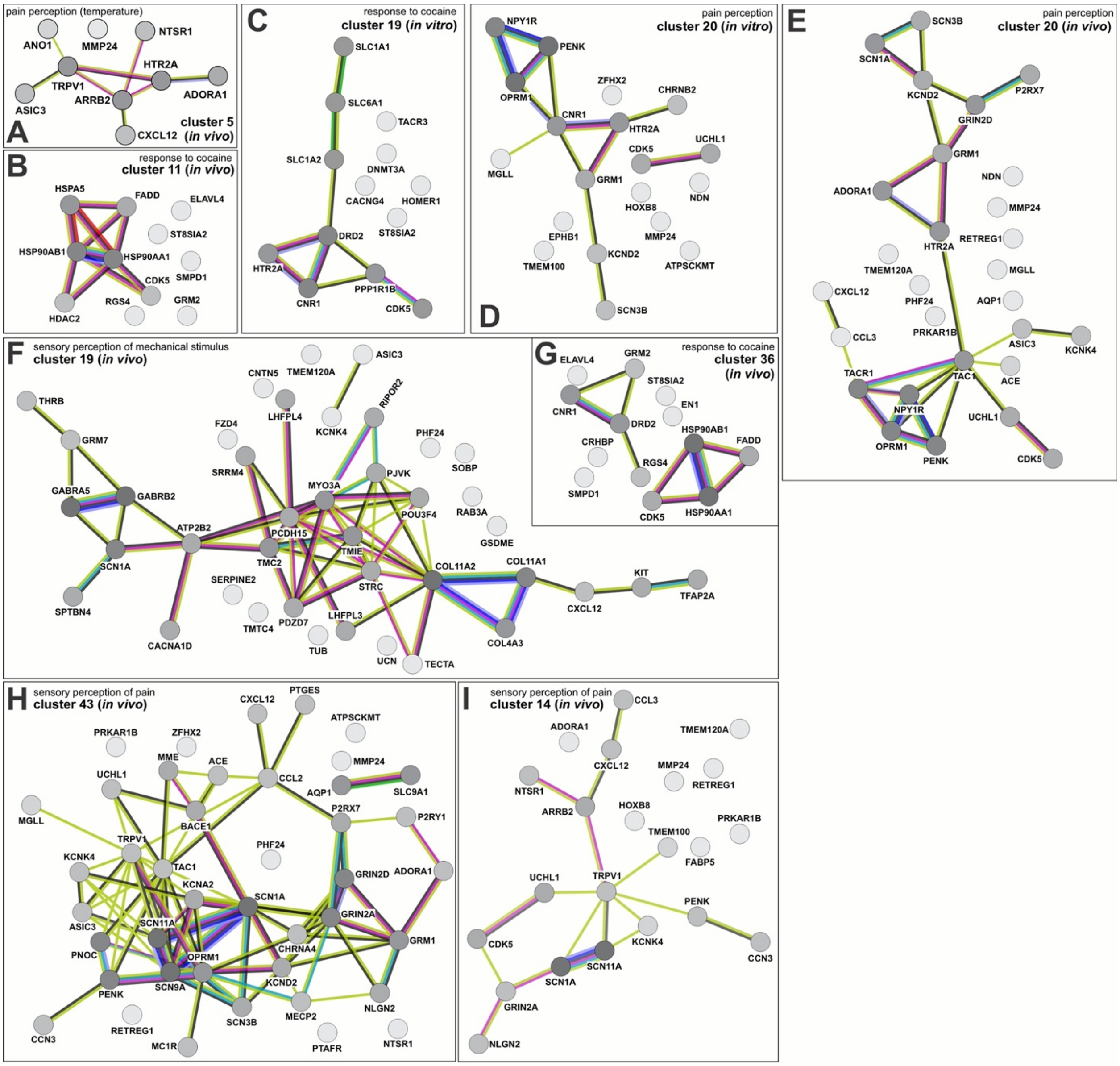


The STRING (search tool for the retrieval of interacting genes/proteins)[59] was used to identify sensory modality networks from genes upregulated in both *in vivo* and *in vitro* dI cell populations. Nodes represent proteins, the lines (or edges) between the nodes represent protein-protein associations. Numerous edges between nodes implies greater evidence (experimentally determined or predicted) for protein-protein associations. STRING further documents evidence of protein associations through the edge color as follows:

Known interactions: experimentally determined (purple) or from curated databases (teal). Predicted interactions: gene neighborhood (green), gene fusions (red), gene co-occurrence (blue). Others: text mining (yellow), co-expression (black), protein homology (periwinkle).

(A, D, E, H, I) Networks associated with pain perception (A) *In-vivo* cluster 5 cells make up the sole network linking temperature with pain, centering around NTSR1, TRPV1, ARRB2 AND HTR2A. Pain circuits regulated by PENK, OPRM1, NPY1R, HTR2A and GRM1 are found in both *in vitro* cluster 20 (D) and *in vivo* cluster 20 (E). The *in vivo* cluster 43 (H) circuit has similarity with *in vivo* cluster 14 (I), sharing the SCN1A, SCN11A, and TRPV1 nodes.

(B, C, G) Protein networks involved in response to cocaine. Both *in vitro* cluster 19 (C) and *in vivo* cluster 36 (G) circuits center around DRD2 and CNR1, further evidence supporting our *in vitro* generated dI4 cells may be functionally similar to their *in vivo* counterparts. (F) Sensory perception of mechanical stimulus protein network stemming from upregulated genes found in *in vivo* cluster 19

**Extended Data Figure 9:** Mechanosensation and balance networks are found among *in vivo* and *in vitro* dI4 clusters

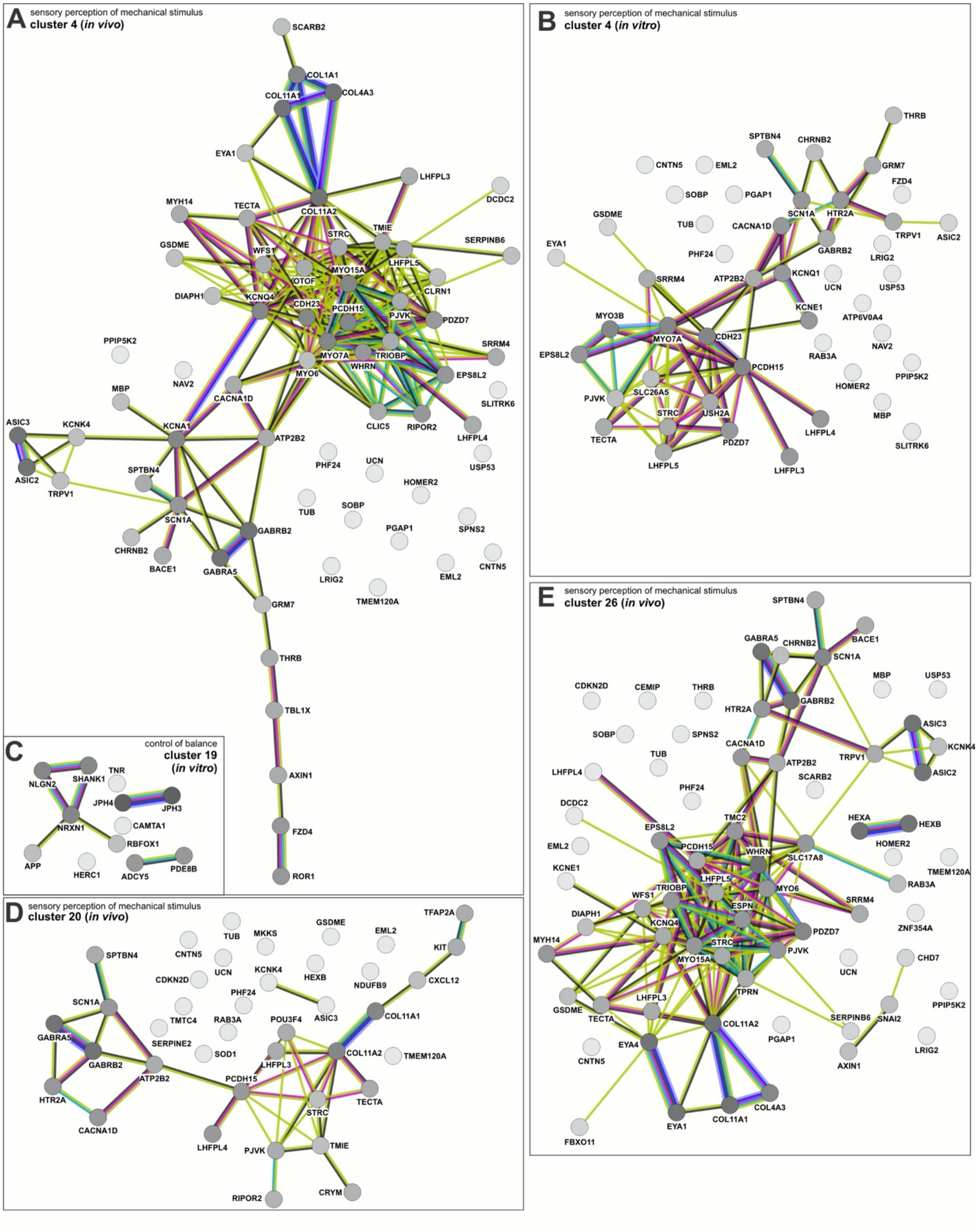

(A, B, D, E) Numerous *in vitro* and *in vivo* clusters contain protein interaction networks involved in mechanosensation. The *in vitro* cluster 4 (B) circuit includes MYO7A, PCDH15, MYO3B and EPS8L2, key nodes also found across *in vivo* clusters 4 (A), 20 (D) and 26 (E).

(C) A protein network associated with balance is found in *in vitro* cluster 19. Notably, this includes SHANK1, NLGN2, and NRXN1, genes with allele variants associated with autism spectrum disorder.

**Extended Data Figure 10:** Pain perception, posture regulation, and mechanosensory networks are found among *in vivo* and *in vitro* dI5 clusters

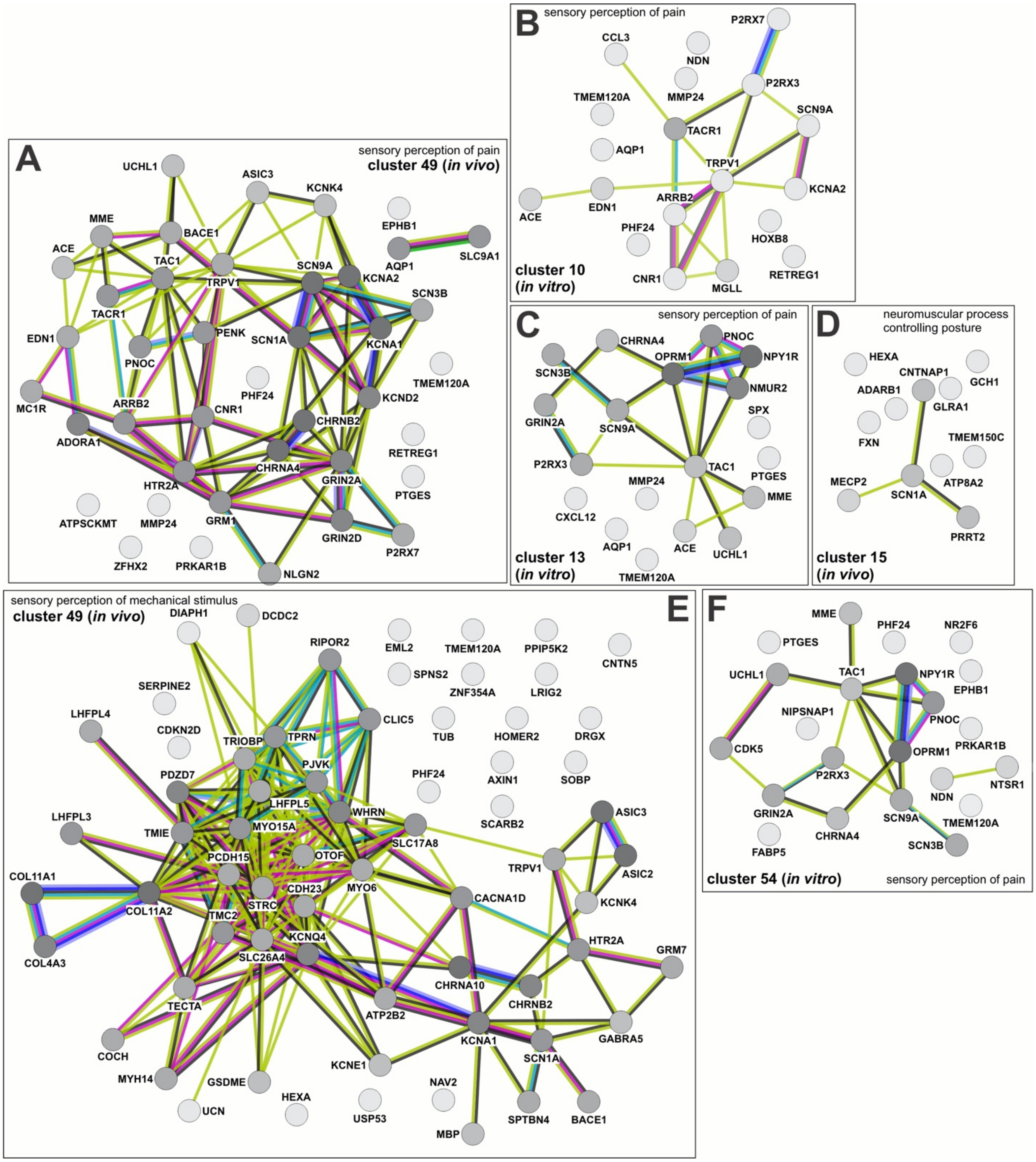

(A-C, F) Networks identified associated with pain perception. The cluster 49 *in vivo* circuit includes strong associations between SCN9A, SCN1A, KCNA1, and SCN3B. Part of this module is recapitulated in *in vitro* cluster 10 (B), with the presence of SCN9A and KCNA2 as well as *in vitro* cluster 54 (F) with the presence of SCN9A and SCN3B. *In vitro* cluster 13 (C) interestingly includes NMUR2, a protein crucial in sensing mechanical itch, which is not present in other identified networks.

(D) Unique to the *in vivo* dataset is a neuromuscular posture regulating network comprised of CNTNAP1, SCN1A, PRRT2 and MECP2.

(E) A large *in vivo* network associated with mechanosensation is found in cluster 49 and includes similar nodes such as PCDH15 described in the clusters in Extended data figure 8 relating to mechanosensation.
